# Single-nuclei isoform RNA sequencing reveals combination patterns of transcript elements across human brain cell types

**DOI:** 10.1101/2021.12.29.474385

**Authors:** Simon A Hardwick, Wen Hu, Anoushka Joglekar, Li Fan, Paul G Collier, Careen Foord, Jennifer Balacco, Natan Belchikov, Julien Jarroux, Andrey Prjibelski, Alla Mikheenko, Wenjie Luo, Teresa A Milner, Lishomwa C Ndhlovu, John Q Trojanowski, Virginia MY Lee, Olivier Fedrigo, Dóra Tombácz, M Elizabeth Ross, Erich Jarvis, Zsolt Boldogkői, Li Gan, Hagen U Tilgner

## Abstract

Single-nuclei RNA-Seq is being widely employed to investigate cell types, especially of human brain and other frozen samples. In contrast to single-cell approaches, however, the majority of single-nuclei RNA counts originate from partially processed RNA leading to intronic cDNAs, thus hindering the investigation of complete isoforms. Here, using microfluidics, PCR-based artifact removal, target enrichment, and long-read sequencing, we developed single-nuclei isoform RNA-sequencing (‘SnISOr-Seq’), and applied it to the analysis of human adult frontal cortex samples. We found that exons associated with autism exhibit coordinated and more cell-type specific inclusion than exons associated with schizophrenia or ALS. We discovered two distinct modes of combination patterns: first, those distinguishing cell types in the human brain. These are enriched in combinations of TSS-exon, exon-polyA site, and distant (non-adjacent) exon pairs. Second, those with all isoform combinations found within one neural cell type, which are enriched in adjacent exon pairs. Furthermore, adjacent exon pairs are predominantly mutually associated, while distant pairs are frequently mutually exclusive. Finally, we observed that human-specific exons are as tightly coordinated as conserved exons, pointing to an efficient evolutionary mechanism underpinning coordination. SnISOr-Seq opens the door to single-nuclei long-read isoform analysis in the human brain, and in any frozen, archived or hard-to-dissociate sample.

## Introduction

Concurrent with the development of single-cell RNA-Seq^1–3^, long-read approaches enabled complete-isoform analysis^4–8^. More recently, long reads empowered the analysis of a few^9,10^ and then thousands of single cells^11,12^ using high-throughput single-cell approaches, including 10x Genomics.

Single-nuclei methods^13–15^ are widely used for many applications and especially for frozen tissues including human brain (Fig. 1a). Single-nuclei datasets contain many partially or fully unspliced RNAs, leading to many reads derived from intronic regions. These reads are usable for gene-count and “RNA-velocity” analyses^16–18^. However, such intronic reads cannot inform on complete isoforms. Another problem for long-read sequencing of 10x Genomics single-nuclei and single-cell libraries are molecules lacking polyA tails, barcodes and Illumina adaptors (Fig. 1b). Such cDNAs are ignored by Illumina machines, but sequenced on Pacific Biosciences (“PacBio”)^19^ and Oxford Nanopore Technologies (“ONT”) platforms, which do not require Illumina adaptors. Here, we present a new approach — <u>s</u>ingle-<u>n</u>uclei <u>iso</u>form <u>R</u>NA <u>seq</u>uencing (termed ‘SnISOr-Seq’) — that overcomes both above problems. Briefly, we employ linear/asymmetric PCR, amplifying cDNAs from the 10x-Genomics partial-read1, near which polyA tails and barcodes reside. This step enriches for polyA-tail- and barcode-containing molecules (Fig. 1c). Second, we use enrichment probes to select cDNA molecules overlapping exons, thereby removing purely intronic molecules (Fig. 1d). We collectively refer to these linear/asymmetric PCR and capture steps as “LAP-CAP”. We then long-read sequence these post-LAP-CAP molecules (Fig. 1e).

**Figure 1:**
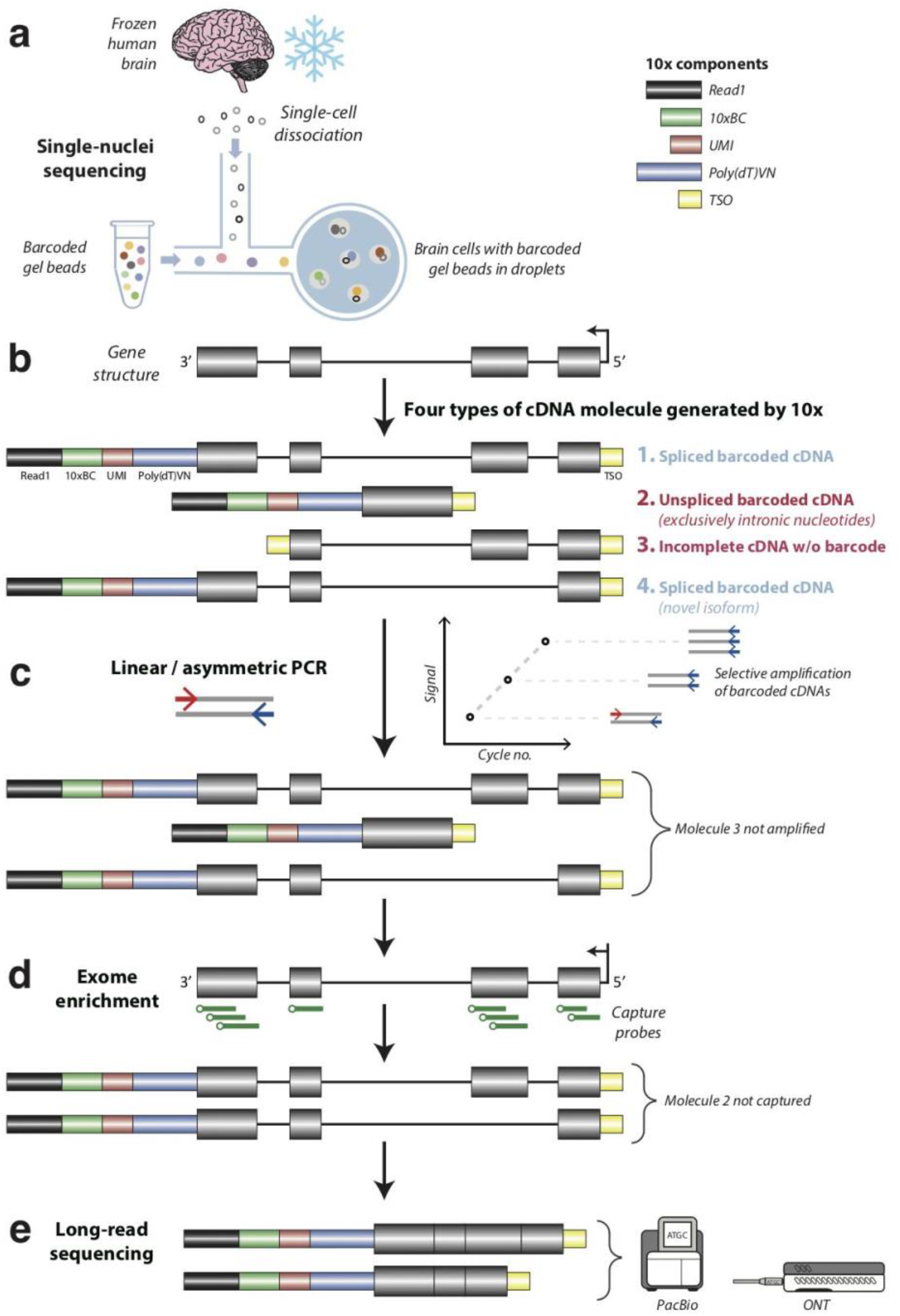
Overview of the single nuclei isoform RNA sequencing (‘SnISOr-Seq’) approach. **a**. The 10x Genomics Chromium platform is used to generate a barcoded cDNA library from a nuclei suspension collected from frozen human brain tissue. **b**. This process generates four types of cDNA molecules: spliced barcoded cDNA (including both known and novel isoforms), unspliced barcoded cDNA (comprising exclusively intronic nucleotides), and incomplete cDNA without a cellular barcode (with TSO on both ends). The latter two molecules compromise efficiency. **c**. Linear/asymmetric PCR is used to selectively amplify barcoded cDNA. **d**. Probe-based exome enrichment step is applied to filter out cDNA molecules which are purely intronic. **e**. Molecules are sequenced on a long-read sequencer (PacBio and ONT). TSO: template-switching oligonucleotide.

Using SnISOr-Seq, we investigate how distinct “RNA variables” – alternative transcription start sites (TSS), exons, and polyadenylation sites (PAS) – are combined into full-length isoforms in the human brain, and to determine the cell-type specific basis of coordination events. We and others have previously investigated the coordination of exon pairs, TSS, and PAS genome-wide^7,20,21^ or specifically for neurexins^22,23^. Mechanisms underlying exon-exon coordination and the influence of promoters on splicing are established^24,25^ for individual genes. Surprisingly, splicing can also influence TSS-choice^26^ and interactions between splicing and 3’-end cleavage have also been described^27^. Likewise, the order of intron-removal from the pre-mRNA has been tackled in yeast^28^. However, how TSS, exon, and PAS combinations specify cell types in the human brain remains unknown, limiting our understanding of brain function. On one extreme, coordination between any RNA-variable pair observed in brain tissue could arise from one or more distinct cell types, i.e. what is observed in bulk brain is also observed in a specific cell-type. On the other extreme, the constitutive use of two RNA variables in one cell type and their skipping in another can underlie the observation of coordination in bulk brain. We find that TSS-exon and exon-PAS coordination follow a similar model to the coordination of distant alternative exons, while adjacent alternative exons follow a different model for cell-type usage. Alternative splicing mis-regulation in disease is established^22,29,30^, however whether these exons are independently affected or hijacked in coordinated units is unknown. Using SnISOr-Seq’s capacity for cell-type specific long-read sequencing, we find that exons associated with distinct diseases exhibit distinct behavior in terms of (i) inclusion variability across cell types and (ii) coordination. Despite common cortical roots, ASD-associated exons show markedly different behavior compared to schizophrenia-, and amyotrophic lateral sclerosis (ALS)-associated exons.

## Results

### An efficient method for removing single-cell artifacts and unspliced RNAs

We first performed single-nuclei 3’-end sequencing of frontal cortex tissue from two healthy donors aged 68 and 61 years from the Penn Brain Bank (henceforth referred to as ‘FCtx1’ and ‘FCtx2’). Employing standard protocols for single-cell analysis^31,32^, we defined 12 clusters representing all major cortical cell types including neurons, astrocytes, oligodendrocytes, microglia, and vascular cells. Among neurons, we observed multiple inhibitory neuron types, including *SST*+, *LAMP5*+, *PVALB*+ interneurons, and layer-specific excitatory neuron types: *RORB+, SEMA3E+, LINC00507+* (Fig. 2a; Supplementary Fig. 1a). We sequenced 8,376 unique molecular identifiers (UMIs) per cell of FCtx1, with excitatory neurons (subtype *RORB* and *SEMA3E*) showing the highest UMI counts per nucleus, and astrocytes and oligodendrocytes the lowest (Supplementary Fig. 2a). These UMI statistics were mirrored by similar gene-per-nucleus trends (Supplementary Fig. 2b). Both FCtx1 and FCtx2 showed high percentages of reads attributed to nuclei and low antisense mappings (Supplementary Fig. 2c). We then used 500 ng of full-length cDNA and performed linear/asymmetric PCR and Agilent exome enrichments (LAP-CAP; Methods) followed by exponential/symmetric PCR. The resulting cDNAs were sequenced on 8 (FCtx1) and 7 (FCtx2) PacBio SMRT cells and 3 (FCtx1) and 2 (FCtx2) ONT PromethION flow-cells. This yielded ∼290×10^6^ long reads with average lengths of 0.9-1.2 kb across technologies and samples (Supplementary Table 1). As a negative control, we sequenced 1 SMRT cell per sample before LAP-CAP. We detected barcodes in long reads as recently published^12,33.^ The barcoded read fraction increased 2.2-fold from naïve single-nuclei long-read sequencing to LAP-CAP (Fig. 2b). Likewise, on-target reads increased 3.3-fold after LAP-CAP, offering additional benefit to the improvements in barcode detection (Fig. 2b). We observed strong correlation in gene expression between FCtx1 and FCtx2 (r=0.947; Fig. 2c), demonstrating SnISOr-Seq’s replicability. Furthermore, the correlation observed for FCtx1 pre- and post-LAP-CAP was relatively strong (r=0.881; Fig. 2d).

**Figure 2:**
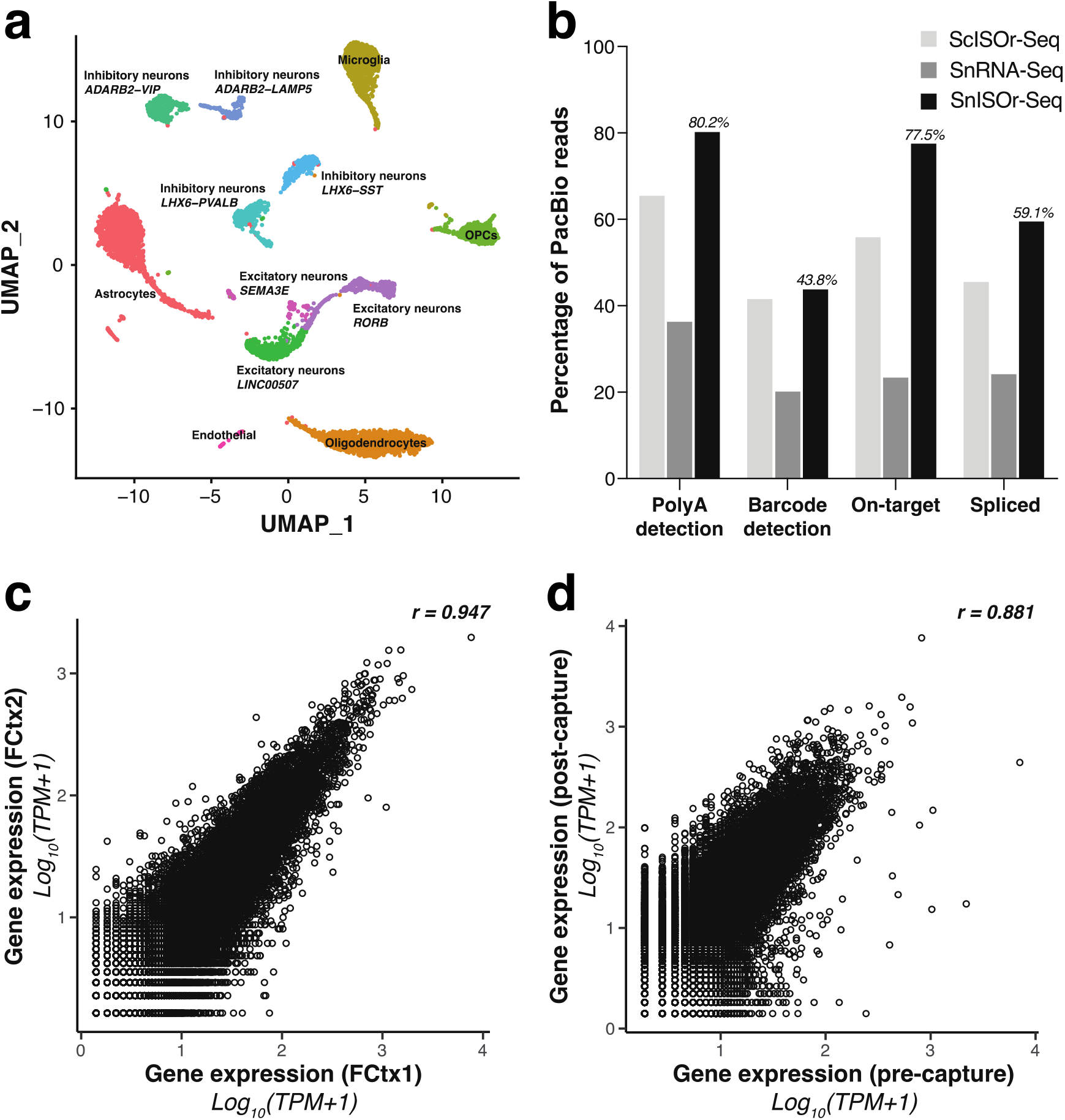
Cell type clustering and enrichment efficiency. **a**. UMAP plot of the FCtx1 sample with each point representing a single nucleus, and colors indicating cell type. **b**. Barplot showing the percentage of PacBio reads with polyA-tails detected, barcodes detected, on-target rate for LAP-CAP, and percentage of spliced reads. Color of the bars indicates the approach used for the comparative analysis. ‘ScISOr-Seq’ refers to single-cell isoform RNA-sequencing, our previously published method^33^; ‘SnRNA-Seq’ refers to regular long-read single-nuclei sequencing, i.e. FCtx1 sample prior to LAP-CAP. **c**. Dotplot of the correlation in PacBio long-read gene expression (log_10_ TPM+1) between FCtx1 and FCtx2. Pearson correlation (r) is indicated. **d**. Dotplot of the correlation in PacBio long-read gene expression for FCtx1 before and after LAP-CAP. TPM: transcripts per million.

For the ONT datasets, we found a median of 515 and 384 reads per nucleus for FCtx1 and FCtx2, respectively (Supplementary Fig. 3a,b). The four major cell types represented 77.9% (FCtx1) and 82.6% (FCtx2) of nuclei (Supplementary Fig. 3c,d), and excitatory neurons consistently had the most reads, UMIs and genes per nucleus (Supplementary Fig. 3e-j). Of note, excitatory neurons had higher counts in FCtx2, mostly at the expense of oligodendrocytes and astrocytes (Supplementary Fig. 3). We observed similar trends for the PacBio libraries (data not shown).

### Single-exon patterns reveal variable inclusion across cell types, including for ASD-associated exons

Microexons are in majority highly included in neurons^34.^ Using alternative exons (Methods) whose genes are expressed in excitatory neurons, inhibitory neurons, astrocytes, and oligodendrocytes, we calculated *Ψ* (percent spliced-in) values for these exons and considered the maximal **Δ***Ψ* (Methods) between these cell types for FCtx1 (Fig. 3a). Consistent with previous observations, the most variable exons were enriched in microexons, defined as ≤27 nt. However, highly variable exons were also enriched for exons ≤54 nt, i.e. twice the maximal length for microexons, and, albeit less pronounced, for ≤75 nt (Fig. 3b). Thus cell-type specific exon inclusion separates shorter exons from longer ones, although far beyond the strict microexon definition. Cell-type specific inclusion of disease-associated exons can pinpoint disease-implicated cell types. We therefore investigated published exons associated with schizophrenia^35^, autism spectrum disorder (ASD)^34,36,37^, and amyotrophic lateral sclerosis (ALS)^38^ for inclusion variability across cell-types. Separating our 5,855 alternative exons into schizophrenia-associated and non-schizophrenia-associated exons, we found no significance (two-sided Wilcoxon-rank-sum test, *p*=0.13) and only a 1.2-fold ratio between the two medians. Likewise, considering ALS, we found a fold change of ratio close to 1, albeit with a significant p-value in one replicate. Thus, the schizophrenia-associated exons used here behave largely like random alternative exons in terms of cell-type specific inclusion. ASD-associated exons, however, behaved differently. ASD-associated exons were dramatically more variable across cell types (two-sided Wilcoxon-rank-sum test, *p*<2.22×10^−16^) with a 2.2-fold higher median than non-ASD-associated alternative exons. To control for previous observations regarding microexons in ASD, we excluded microexons from our comparative analysis, and the observation remained true (Fig. 3c). Importantly, this variability of ASD-associated exons does not stem from inclusion in one specific cell-type overall. Indeed, apart from many exons highly included in all four cell types, we observed two other groups: one group exhibits high inclusion in excitatory and inhibitory neurons but low inclusion in astrocytes and oligodendrocytes. Conversely, a second group showed high inclusion in astrocytes and oligodendrocytes but low inclusion in both neuronal types. More complicated cell-type specific arrangements were also observed, albeit less often (Fig. 3d).

**Figure 3:**
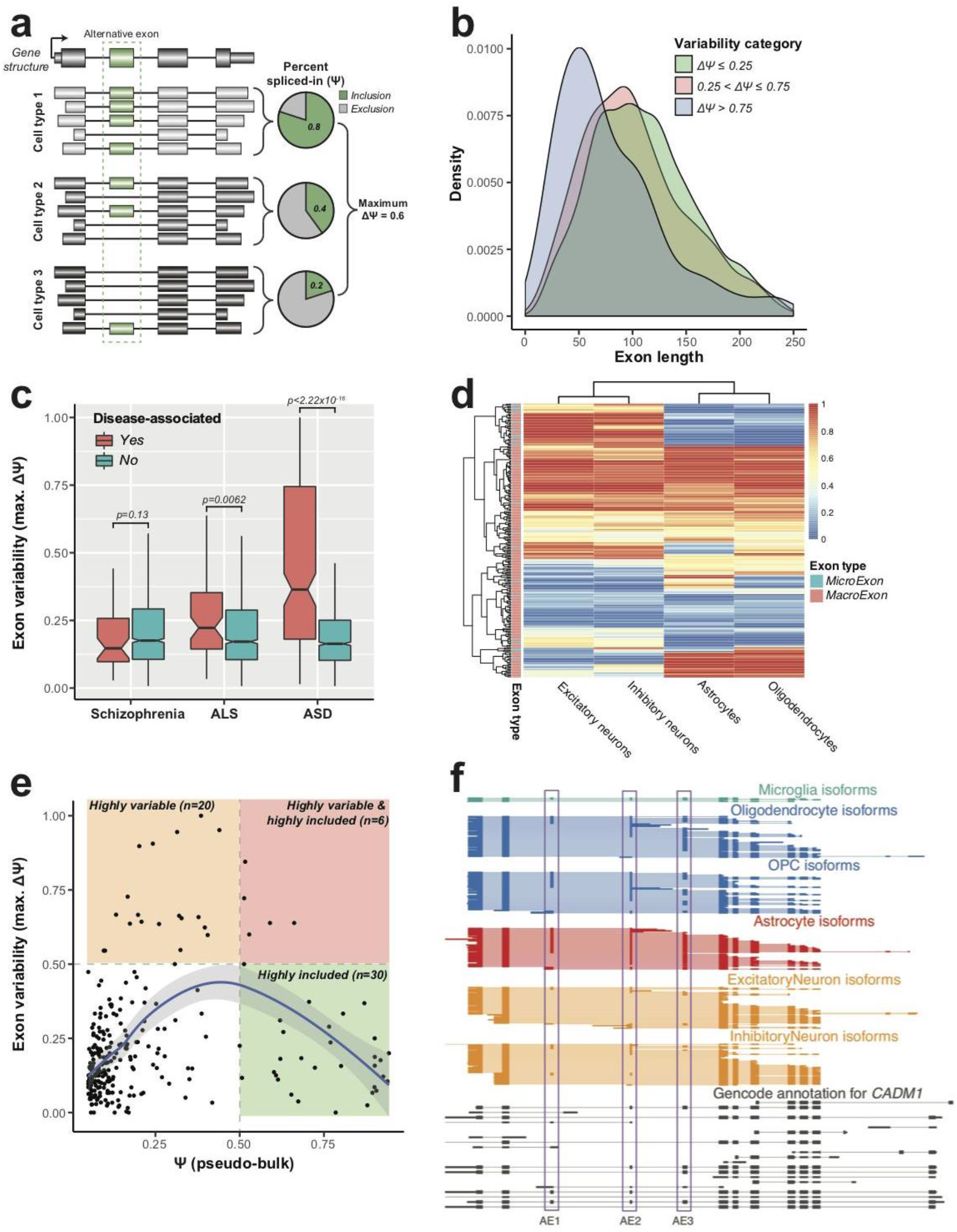
Alternative usage of single exons. **a**. Schematic illustrating how percent spliced-in (*Ψ*) is calculated for an alternative exon (green). The exon shows different levels of inclusion across three cell types, with variability defined by the max*Ψ* - min*Ψ*. **b**. Density plot of the variability in exon inclusion across the four major cell types, and exon length on the X-axis. Colors indicate the discrete categories that the variability values have been divided into. **c**. Boxplots of the variability of exon inclusion for alternative disease-associated exons (red) compared to alternative exons with no known association with that disease (green). P-values obtained from a two-sided Wilcoxon rank sum test. Investigated diseases represented on the X-axis. **d**. Heatmap of the exon inclusion level for ASD-associated exons where each row is an exon and each column is a cell type. Annotation of exon classification as a microexon (≤27 bp) and macroexon (>27 bp) on the left. Color of each cell in the heatmap indicates *Ψ* value. **e**. A dotplot of the *Ψ* of pseudo-bulk (i.e. across all nuclei) on the X-axis and exon variability (max*Ψ* - min*Ψ*) across cell types on the Y-axis. This plot includes all novel exons that had ≥10 reads in ≥2 cell-types. Regression curve with confidence interval obtained using the loess fit. Boundaries of low and high variability and inclusion defined at 0.5 on both axes. **f**. Full-length transcript expression broken down by cell type for the *CADM1* gene. Each horizontal line indicates one transcript, colored by cell type of origin, while clustered blocks indicate exons. Black transcripts denote annotated GENCODE transcripts. Purple boxes highlight three alternative exons labeled AE1-AE3. ALS: amyotrophic lateral sclerosis; ASD: autism spectrum disorder.

Of the above 5,855 alternative exons, 586 were novel with respect to the GENCODE annotation (v34) and had ≥10 overlapping reads in ≥1 cell type. The question of which exons should be included in state-of-the-art annotations is of high importance^39^. Obviously, exons with high bulk-tissue *Ψ* should be included in annotations and likewise exons that are variably included across cell types. We therefore calculated an overall *Ψ* from all nuclei combined (termed “pseudo-bulk”) and the maximal **Δ***Ψ* between the cell types. Thus, 20 non-annotated exons should be included in annotations, because of high variability (**Δ***Ψ*≥0.5) across cell types, implying at least one cell type with high inclusion, despite low overall inclusion (pseudo-bulk *Ψ* ≤0.5). A further 6 novel alternative exons should be included in annotations, as they had both high pseudo-bulk inclusion (*Ψ*>0.5) and high variability (**Δ***Ψ*≥0.5). This observation implies at least one cell type, in which these exons are mostly skipped. Finally, another 30 exons showed high pseudo-bulk inclusion (*Ψ*≥0.5) and low variability (all **Δ***Ψ*s≤0.5), indicating their preferential inclusion in most cell types. The remaining 206 novel alternative exons were lowly included (pseudo-bulk *Ψ*≤0.5) and showed low variability (all **Δ***Ψ*≤0.5). While the arbitrary 0.5 cutoffs can be debated, the combined use of pseudo-bulk inclusion and *Ψ* variability helps characterize an exon’s importance (Fig. 3e). The above observations were broadly replicable in FCtx2 (Supplementary Fig. 4). *CADM1* illustrates multiple highly cell-type specific alternative exons in one gene (Fig. 3f). Three alternative exons are more highly included in astrocytes, oligodendrocyte precursor cells (OPCs) and oligodendrocytes than in both excitatory and inhibitory neurons. Two alternative exons (labeled “AE2” and “AE3”) are very highly included in astrocytes. The ASD association of AE3 in glial cell types, suggests exploring its possible glial mis-regulation in ASD.

### Combinations of exon, TSS, and PAS show distinct pairing rules

We and others have investigated patterns of exon combinations, however the frequency of different combination patterns remains unclear. Two exons may be paired non-randomly (i.e., in a coordinated fashion) or randomly. The former can represent a tendency for mutual inclusion or exclusion (Fig. 4a). We first considered alternative exon pairs.. After statistical testing and false-discovery rate (FDR) calculation, only one exon pair per gene was retained to avoid the discovery of patterns representing few genes with many exon pairs (Methods). Among neighboring exon pairs, 71.4% of tested pairs, each represented by a 2×2 table, showed a significant association at FDR=0.05 and an absolute-value log-odds ratio (LOR) ≥1. By definition, this fraction decreases for higher log-odds ratios. However, even for an |LOR|≥7, a 128-fold enrichment of two of the exon combinations over the other two, >=50% of exon pairs showed non-random pairing (Fig. 4b). For distant alternative exon pairs — i.e., those with intervening exons, which we investigated before^7,12,20^ — this fraction was drastically lower (Fig. 4c). An example of neighboring coordinated exons is the *WDR49* gene. Two neighboring coding exons are positively and perfectly coordinated, i.e., all molecules include either both exons or none, while molecules with only one exon are not observed. In this case, coordination of both exons originates from an individual cell type, namely astrocytes (Fig. 4d). Of note, the above gene fractions harboring a coordinated exon pair (see Fig. 4b,c) represent lower bounds, as deeper sequencing could turn non-significant pairs significant. Importantly, adjacent coordinated alternative exons showed stronger coordination than distant coordinated exon pairs (Fig. 4e). Furthermore, distant exon pairs frequently show mutual-exclusion coordination, i.e., a negative LOR, whereas this is dramatically less likely for adjacent exon pairs (Fig. 4f) which dominate our dataset. Similar observations arise for FCtx2 (Supplementary Fig. 5a-d). Consistent with adjacent mutually exclusive exons often exhibiting sequence homology^40^ and given that our adjacent coordinated exons are mostly mutually inclusive, we find almost no sequence similarity between these exon pairs. Given their tight coordination, we hypothesized that coordinated adjacent exon pairs would be highly evolutionarily conserved. Surprisingly, however, we observe very low non-significant correlation (Pearson R=0.089, *p*=0.063; Spearman-rank-sum correlation=0.0526; *p*=0.274) between phastCons scores^41^ of the less conserved exon and the overall coordination strength. Thus, evolutionarily recent exons are almost as tightly paired with other exons as ancient exons, pointing to an efficient evolutionary mechanism achieving coordination (Fig. 4g). The above correlation between phastCons scores and exon-pair coordination was low, but significantly positive. However, for coordination events between PASs and alternative exons, the correlation was negative – suggesting a different mechanism that coordinates PAS with alternative exons (Fig. 4h).

**Figure 4:**
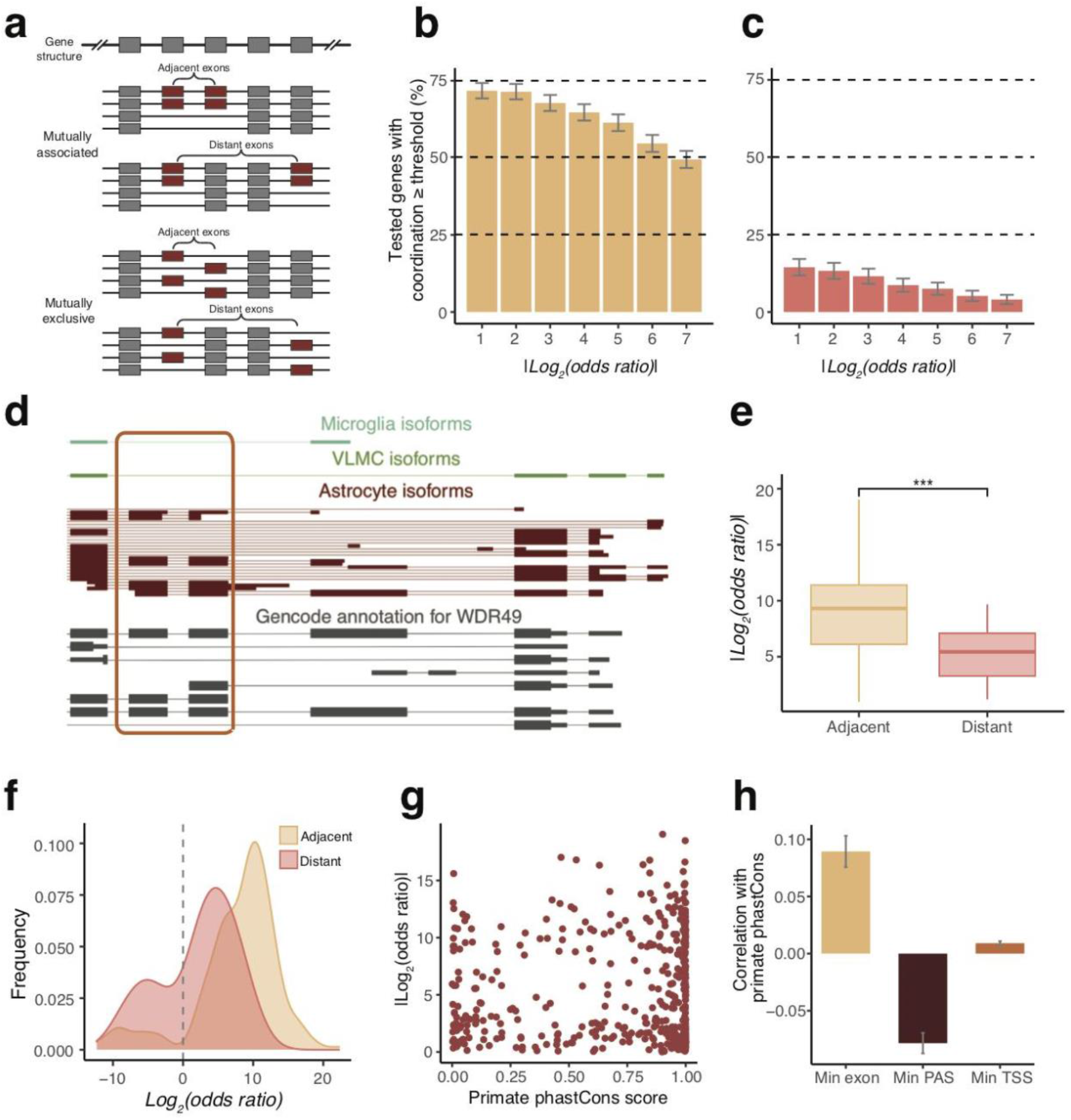
Coordination of adjacent and distant exon pairs. **a**. Schematic showing the types of exon coordination patterns when considering two alternative exons (red). Mutual inclusion of distant and adjacent exons (top) and mutual exclusion of distant and adjacent alternative exons (bottom). **b-c**. Barplot showing percent of tested genes in pseudo-bulk with significant exon coordination (FDR≤0.05) for adjacent (b) and distant (c) exon pairs at various log-odds ratio (LOR) cutoffs on the X-axis. **d**. Zoomed-in region of adjacently coordinated exons for the *WDR49* gene. Each horizontal line indicates one transcript, colored by cell type of origin, while clustered blocks indicate exons. Grey transcripts denote annotated GENCODE transcripts. Orange box highlights the coordinated exons, the transcripts of which make up a majority of the pseudo-bulk expression. **e**. Boxplots of the |LOR| for significant genes (FDR ≤ 0.05) on the Y-axis plotted against adjacent and distant exon pairs seen in (b) and (c) on the X-axis. p-value obtained from two-sided Wilcoxon rank sum test. **f**. Density plot for the LOR for adjacent and distant exon pairs. **g**. Dotplot of the absolute value of the LOR of coordination for tested exon pairs on the Y-axis with the minimum primate phastCons score from the exon pair on the X-axis. **h**. Barplot of the pearson correlation value of the phastCons score on the Y-axis grouped by the minimum of two tested exon pairs, minimum score among the TSS associated with a given exon, and minimum score among the PAS associated with a given exon. Each group denoted separately on the X-axis. Error bars (b,c,h) indicate standard error of the point estimate.

### Coordination of exon pairs observed in bulk mostly stems from coordination in specific cell types

We then examined whether the coordination patterns at pseudo-bulk level were detected in at least one high-level cell type, or whether they represent a heterogeneous mixture of homogeneous cell-type specific patterns. Here, we considered excitatory neurons, inhibitory neurons, astrocytes, oligodendrocytes and OPCs as high-level cell types. Among the mostly adjacent coordinated pseudo-bulk exon pairs testable in ≥1 cell type, 89% were significantly coordinated in ≥1 cell type. More precisely, 41.7% were coordinated in one cell type, 21.3% in two, and 24% in three, four or five cell types (Fig. 5a). These observations were broadly conserved in FCtx2 (Supplementary Fig. 6a). In all five cell types investigated, ≥50% of testable (mostly adjacent) exon pairs showed significant coordination, but percentages varied between cell types. Indeed, for astrocytes, only 54.08%, while for oligodendrocytes and OPCs 67.14% and 72.72% showed coordination respectively (Fig. 5b; Supplementary Fig. 6b). Importantly, two distinct models can explain why an exon pair that is testable in pseudo-bulk is not testable in a cell type. First, read counts in a cell type, which are by definition lower than or equal to those in the pseudo-bulk, may simply be too low to allow for **χ**^2^ testing – a model purely technical in nature. Second, one of the exons may become constitutively included or skipped in the cell type (Methods); this implies that the **χ**^2^ criterion for testability is violated - a model biological in nature. Importantly, distant alternative exon pairs are ∼2-fold more likely to have ≥1 exon constitutively included/skipped in ≥1 cell type than adjacent alternative exons (Fig. 5c). This finding was replicated in each cell type separately, although non-overlapping 95%-confidence intervals were only observed in excitatory neurons, inhibitory neurons and oligodendrocytes (Fig. 5d).

**Figure 5:**
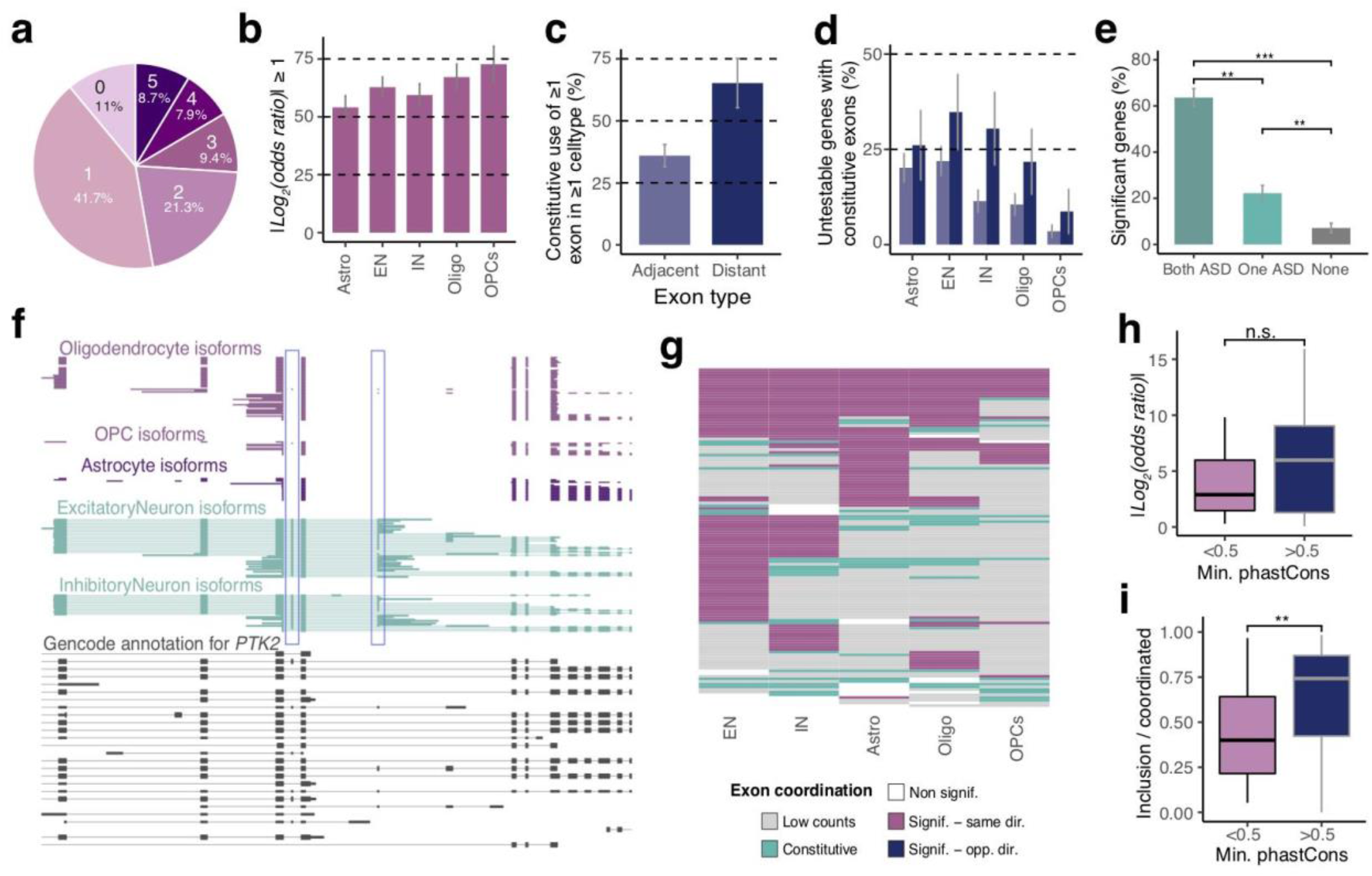
Exon coordination patterns are observable across multiple cell types. **a**. Pie chart indicating the number of cell types where an exon pair is significant (FDR ≤ 0.05 and abs(LOR) ≥1) given that the exon-pair is testable in at least one cell type, and is significant when tested in pseudo-bulk. **b**. Barplot indicating percentage of tested exon-pairs (one per gene) that were deemed significantly coordinated (FDR ≤ 0.05 and abs(LOR) ≥ 1). Breakdown by cell type on the X-axis, given significance in pseudo-bulk. **c**. Barplots denoting percentage of genes which are not testable in any cell-types because at least one exon became constitutively included or excluded in at least one cell type. Values split by whether the exon pair is adjacent or distant and denoted on the X-axis. **d**. Barplots denoting percentage of genes which are not testable in specific cell types because at least one exon became constitutively included or excluded in that cell type, and colored by whether the exon pairs are adjacent or distant. **e**. Barplots showing percent of tested genes that are significant in pseudo-bulk for distant exons pairs where either one, or both, or neither of the exons are known to be associated with autism spectrum disorder (ASD). p-values obtained from two-sided Fisher’s exact test. **f**. Zoomed-in region of distantly coordinated exons for the *PTK2* gene. Each horizontal line indicates one transcript, colored by cell type of origin, while clustered blocks indicate exons. Grey transcripts denote annotated GENCODE transcripts. Blue boxes highlight the coordinated exons. **g**. Heatmap showing cell types as columns and exon pairs that were testable in at least one cell type as rows. Each element of the heatmap is colored by whether the exon-pair showed significant coordination and the direction of coordination with respect to the pseudo-bulk (same: pink or opposite: blue), was not significant (white), was not testable in a particular cell type because it had too few counts to satisfy the **χ**^2^ criterion (grey) or because one or both exons became constitutively included in a particular cell type (teal). **h**. Boxplots of the absolute value of the LOR of excitatory neuron reads for tested exon pairs denoted on the Y-axis and split by whether the minimum phastCons score of the two exons was greater than or less than 0.5. Significance calculated using two-sided Wilcoxon rank sum test. **i**. Boxplots for excitatory neuron reads showing the double inclusion level of an exon pair divided by the number of reads that were coordinated. X-axis indicated whether the minimum phastCons score of the two exons was greater than or less than 0.5. Significance was calculated using the two-sided Wilcoxon rank sum test. Error bars (b-e) represent the standard error of the point estimate

In addition to, and partially based on, our previous observation of ASD-associated exons being more variably spliced than others, we also find that pairs of ASD-related exons are highly coordinated. Indeed, in 63.6% of cases where both exons of a distant exon pair were linked to ASD, the two exons were coordinated. When neither exon was ASD-associated, only 7.14% of exon pairs were coordinated (two-sided-Fisher’s exact test, *p*=2.96e-5 comparing to both exons being ASD-associated; *p*=0.023 comparing to one ASD-associated exon; Fig. 5e, Supplementary Fig. 6c). An example of distant coordinated ASD-associated exons with a strong cell-type specific component is the *PTK2* gene, which encodes for FAK and influences axonal growth regulation and neuronal cell migration^42^. Two alternative microexons of 18 and 21 bp are highly included in excitatory and inhibitory neurons (co-inclusion score = 0.8 and 0.7; Methods), but are almost completely skipped in glial types (co-inclusion score = 0, 0.02, and 0 respectively for astrocytes, oligodendrocytes, and OPCs). Thus, both exons exhibit very high variability among cell types (Fig. 5f). This further motivates single-cell long-read investigations of ASD.

All significant cell-type specific exon coordination values pointed in the same direction as in the pseudo-bulk. That is, coordination values for adjacent exon pairs observed in bulk reflect coordination in ≥1 cell type. Excitatory neurons, inhibitory neurons and astrocytes clearly recapitulated more coordination events from the pseudo-bulk than oligodendrocytes and OPCs – likely owing to their higher nuclei numbers. Additionally, because of the strong tendency for mutual inclusion for adjacent exons, the majority of molecules represent the both-included and the both-skipped isoforms at the expense of including only one exon (Fig. 5g). In FCtx2, excitatory neurons dominated the genes that were significantly coordinated in bulk due to high excitatory neuron number in FCtx2 (Supplementary Fig. 6d).

Consistent with our pseudo-bulk observations, we found no significant association between exon conservation and coordination at cell-type level (excitatory neurons depicted as a representative cell type here, Fig. 5h). In contrast to this observation, conservation was significantly associated with the inclusion of both alternative exons – at the expense of the skipping of both exons, an observation replicable in FCtx2 (Fig. 5i; Supplementary Fig. 6e-f).

### TSS-exon and exon-PAS coordination often stems from constitutive use of variable sites in distinct cell types

When tracing coordinated exon-TSS events into five major cell types, we observed drastically different behavior compared to that of adjacent exon pairs: in 71% of cases, significant coordination was not observed in any cell type, while in ∼24% and 4.9% coordination was found in one and two cell types respectively. Significance in ≥3 cell types, however, was never observed, and the overall proportion of genes exhibiting TSS-exon coordination were less than 10% at all investigated ***ΔΠ*** cutoffs (Fig. 6a-b; Methods). Contrarily to adjacent exon pairs, constitutive use of one alternative site (TSS or exon) in a cell type occurred frequently - and broadly consistently in all five cell types (Fig. 6c, teal). This is exemplified by the *TBCB* gene, with two main TSS. When transcription is driven from the upstream TSS, the first acceptor (“Acc1”; Fig. 6d, left) is used, while a downstream TSS’s usage mostly results in straight unspliced transcription over Acc1, although the removal of the short 111 nt intron (“Intron2”) is possible. These observations are apparent in the pseudo-bulk and in astrocytes. In excitatory and inhibitory neurons however, the upstream TSS is nearly constitutive. Thus, coordination or coordination testing using **χ**^2^ statistics is impossible. In summary, the TSS-exon coordination observed in the pseudo-bulk is present in astrocytes but not in neuronal cell types (Fig. 6d, left). For the *TBCB* gene, reads selected for 5’ completeness also showed a remarkable pattern of being 3’ complete (Fig. 6d, right). Exon-PAS pairs were overall more consistent with the exon-TSS pairs than with exon-exon pairs in terms of how many individual cell types a coordination event was observed in, and the results were consistent in FCtx2 (Fig. 6e-f; Supplementary Fig. 7a-d). Likewise, constitutive inclusion/skipping of either the exon or PAS in a cell type was observed far more often than for exon-exon pairs and slightly less than for exon-TSS pairs (Fig. 6g; compare this result with Fig. 6c and 5g).

**Figure 6:**
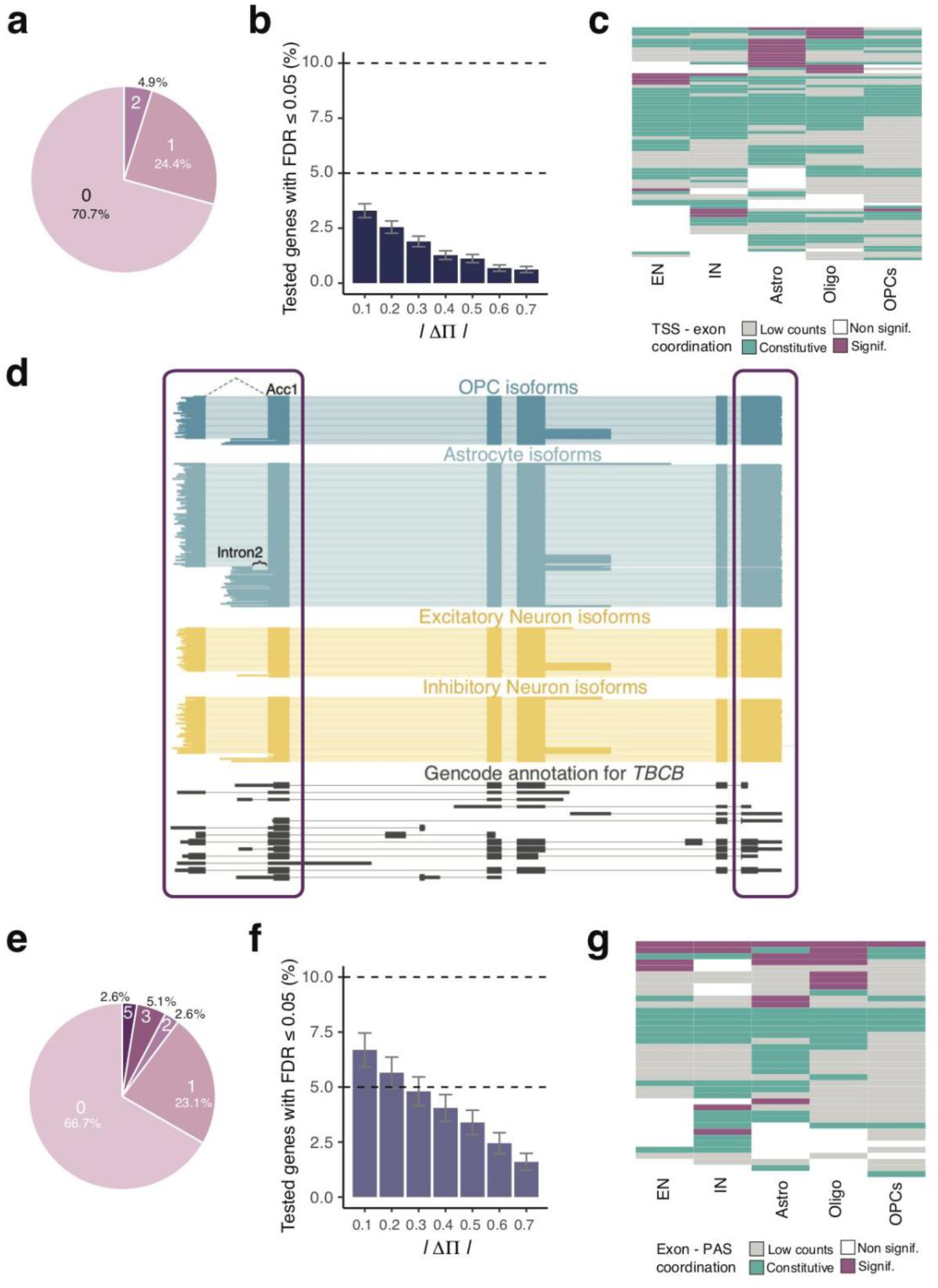
Exon - end site coordination is distinctly mediated by individual cell types. **a**. Pie chart indicating the number of cell types where an exon-TSS pair is significant (FDR ≤ 0.05 and ***ΔΠ*** ≥1) given that the exon-TSS pair is testable in at least one cell type, and is significant when tested in pseudo-bulk. **b**. Barplot showing percent of tested genes in pseudo-bulk with significant exon-TSS coordination on the Y-axis and various ***ΔΠ*** cutoffs on the X-axis. **c**. Heatmap showing cell types as columns and exon-TSS pairs that were testable in at least one cell type as rows. Each element of the heatmap is colored by whether the exon-TSS pair showed significant coordination (pink), was not significant (white), was not testable in a particular cell type because it had too few counts to satisfy the **χ**^2^ criterion (grey), or because the exon or one or more TSS became constitutively included in a particular cell type (teal). **d**. Full-length transcript expression broken down by cell type for the *TBCB* gene. Each horizontal line indicates one transcript, colored by cell type of origin, while clustered blocks indicate exons. Grey transcripts denote annotated GENCODE transcripts. Purple box highlights the region of interest. **e**. Pie chart indicating the number of cell types where an exon-PAS pair is significant given the same testing conditions as in (a). **f**. Barplot showing percent of tested genes in pseudo-bulk with significant exon-PAS coordination on the Y-axis and various ***ΔΠ*** cutoffs on the X-axis. **g**. Heatmap showing cell types as columns and exon-PAS pairs that were testable in at least one cell type as rows. Each element of the heatmap is colored as in (c). Error bars (b,f) indicate standard error of the point estimate.

## Discussion

Elucidating combination patterns of TSSs, exons, and PAS is necessary for a comprehensive understanding of biology, since these patterns define full-length isoforms carrying protein-coding information. Understanding the cell-type specific combination patterns in healthy tissue, particularly of disease-associated exons, is paramount to uncovering affected patterns in disease. Moreover, brain-region and cell-type specific isoform expression may be critical to understanding the clinical relevance of potentially deleterious variants of uncertain significance observed in patient genomes. To investigate these questions we have developed single-nuclei isoform RNA sequencing (SnISOr-Seq, Fig. 1), an approach applicable to any single-nuclei RNA-seq library. While single-nuclei RNA-seq is employed for many tissues, it is especially important for frozen samples, for which entire cell isolation is difficult, the most prominent example being the human brain.

SnISOr-Seq reveals cell-type specific inclusion of individual exons and much more. Consistent with previous reports, we found that microexons (i.e. exons ≤27 bp) show more variable inclusion across cell types than longer exons. However, exons significantly longer than 27 bp (up to ∼75 bp) also show high variability. Of course, the exact cutoff may vary with the definition of “highly-variable” and “non-variable” exons. Importantly, ASD-associated exons, even when excluding microexons, show higher inclusion variability across the four major cell types than random alternative exons. In contrast, the trend for schizophrenia- or ALS-associated exons is dramatically weaker or non-existent. In other words, although a fraction of ASD-associated exons have similar inclusion in the four major cell types, a greater proportion than for other diseases show cell-type specificity. Some ASD-associated exons are highly included in astrocytes and oligodendrocytes while others are more frequently used in neurons. The presence of both cell-type biased and unbiased patterns implies that these exons are not well-investigated by FACS-sorting a single cell type. Single-cell investigations of exon pairing may be even more important for schizophrenia and ALS-associated exons, which are no more variably included across cell types than background exons. These observations raise the fundamental question: whether in disease, the inclusion of these disease-associated exons are altered in all cell types equally, or whether their *Ψ* values change in particular cell types. While splice sites are fundamentally important for understanding the transcriptome, the combination patterns of variable sites inform (i) whether one variable influences another variable’s definition, and (ii) how function might be encoded on individual RNA molecules.

Ample research has investigated exon pairs, TSS-exon pairs, as well as exon-PAS pairs. However, until now, a comparative analysis of these had been lacking in human brain. We find that adjacent exon pairs are combined more often and less randomly than distant pairs. In fact, the majority of genes tested showed coordination of ≥1 adjacent exon pair. Importantly, the gene fraction with coordinated exons could increase even further with deeper sequencing. Moreover, adjacent coordinated alternative exons are almost always mutually inclusive, while distant alternative exons exhibit more mutually exclusivity. TSS-exon pairing and exon-PAS pairing show comparable coordination to distant alternative exons - but significantly less than adjacent exon pairs.

Considering cell-type specific RNA expression, we surprisingly find that three types of coordination, namely TSS-exon, exon-PAS, and distant exon-exon coordination follow the same rule: these types of coordination are often observed at the pseudo-bulk level but cannot be traced into distinct human brain cell types. Rather, they arise by one combination being expressed in one cell type and a different combination occurring in another. Thus, these types of coordination most often reflect the diversity of isoform expression distinguishing cell types (Fig. 7a-c). Adjacent alternative exons, however, follow another pattern: whenever read counts suffice to trace coordination into specific cell types, we usually find one, and often multiple, cell types in which this coordination occurs (Fig. 7d).

**Figure 7:**
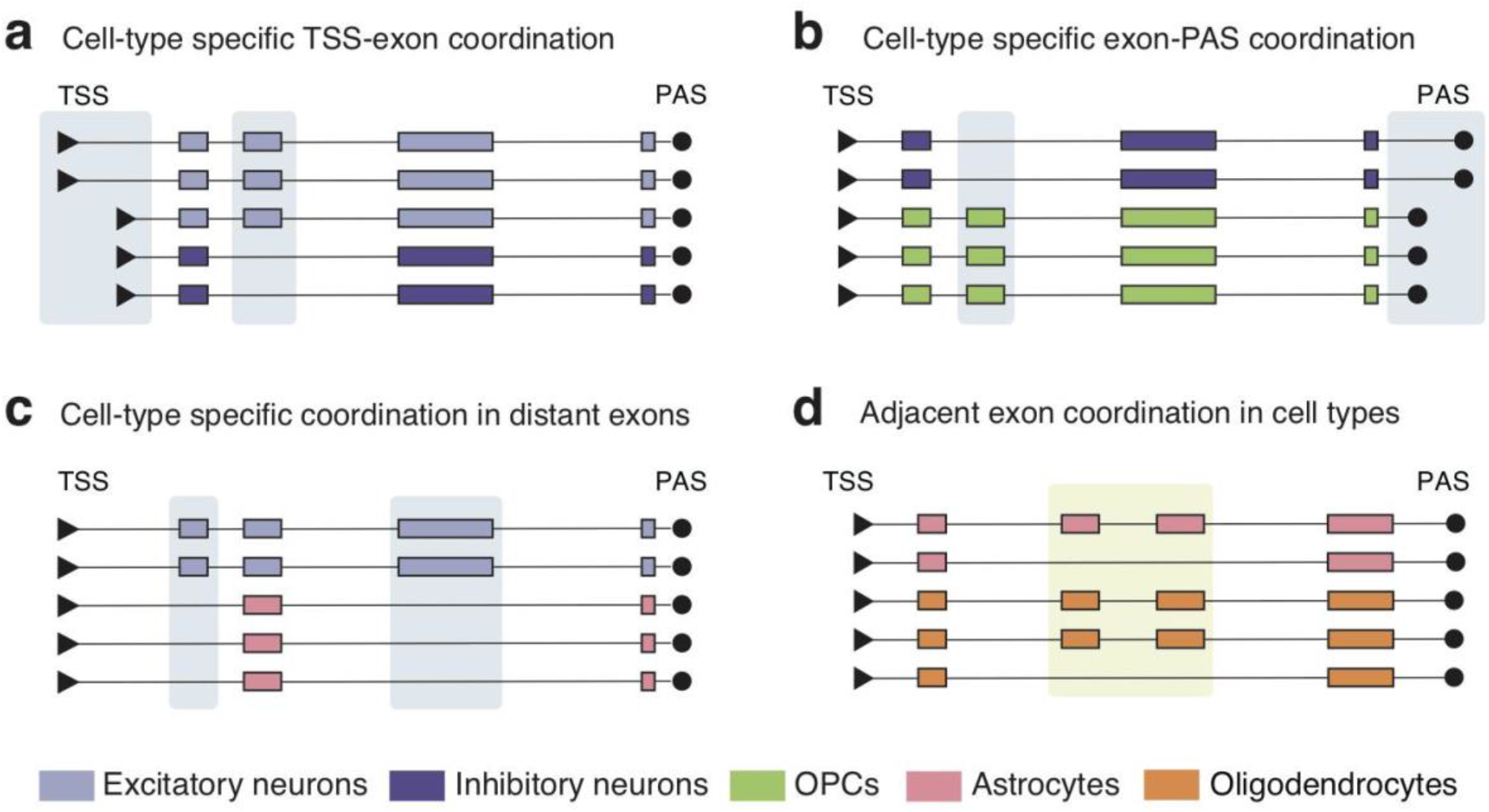
Four different models for cell-type specific coordination of transcriptional elements. **a**. Model for TSS-exon combinations. **b**. Model for exon-PAS combinations. **c**. Model for distant exon combinations, which show a preference for mutual exclusion. **d**. Model for adjacent exon combinations, which show a preference for mutual association. Colors indicate different cell types. TSS: transcription start site; PAS: polyadenylation site.

Thus, when mutual inclusivity vs. exclusivity and cell-type specificity of coordination are considered, TSS-exon, PAS-exon, and distant exon pairs follow one model, while adjacent exon pairs follow a markedly different one. Interestingly, distant alternative exons are highly coordinated and cell-type specific when both exons are associated with ASD. Thus, splicing investigations of the brain in general and a deeper understanding of the role of these exons in neurological disease can benefit from further investigations enabled by SnISOr-seq.

## Acknowledgments

We thank Jason McCormick and Tomas Baumgartner from Weill Cornell Medicine Flow Cytometry Core Facility for FACS assistance, and Dong Xu, Xing Wang, Adrian Tan, and Jenny Xiang from the Genomics Resources Core Facility for performing RNA sequencing. We thank Dr. Christopher Mason for use of his PromethION machine. We also thank Weill Cornell Medicine Scientific Computing Unit (SCU) for use of their computational resources. H.U.T. is supported by NIGMS grant 1R01GM135247-01, Brain Initiative grant 1RF1MH121267-01, NIDA grant U01 DA053625-01, and by the Feil Family Foundation. M.E.R is supported by the National Institute of Health (NIH) grants 1R01NS105477, P01HD067244, U54NS11717, and by the Feil Family Foundation. L.G. is supported by National Institute of Health (NIH) Grants R01AG051390, U54NS100717, R01AG054214, and JPB Foundation. T.A.M. is supported by NIH grants DA08259 and HL136520. J.Q.T. is supported by NIH grant U19 AG062418. E.D.J and O.F. are supported by funds from HHMI. S.A.H. is supported by an Australian NHMRC Early Career Fellowship (APP1156531). A.M. and A.D.P. are supported by St. Petersburg State University, Russia (grant ID PURE 73023672). Computational analysis was performed with the help of the Research park of St. Petersburg State University Computing Center.

## Author Contributions

S.A.H., W.H., A.J., H.U.T. conceived the project and designed experiments. W.H., L.F., P.G.C. performed experiments. S.A.H, W.H., A.J., C.F., N.B., A.P., A.M., J.J., H.U.T. conducted computational analyses. T.A.M., L.C.N., O.F., D.T., M.E.R., E.J., Z.B., L.G., H.U.T. supervised the project. V.M.Y.L., J.Q.T. contributed key reagents. S.A.H., W.H., A.J., H.U.T. wrote the manuscript. All authors participated in the review and editing of the manuscript.

## Methods

### Experimental model and subject details

Two healthy human brain samples (mid-frontal cortex) used for this study were obtained from tissue banks maintained by the Center for Neurodegenerative Disease Research (CNDR) and University of Pennsylvania Alzheimer’s Disease Core Center (ADCC) according to Institutional Review Board-approved protocols. Neither subject had pre-existing neurodegenerative or neurological disease. The postmortem interval was 14 h for FCtx1 (age 68, Male) and 6 h for FCtx2 (age 61, Male). Tissues were flash frozen and kept at -80°C until processing.

### Single nuclei isolation and 10x Genomics 3’ single-nuclei library construction

Single nuclei suspension was isolated from fresh-frozen human brain samples using a protocol adapted from previous studies with modifications^43,44.^

Approximately 30 mg of frozen tissue of each sample was dissected in a sterile dish on dry ice and transferred to a 2 mL glass tube containing 1.5 mL nuclei pure lysis buffer (MilliporeSigma, catalog no. L9286) on ice. Tissue was completely minced and homogenized to nuclei suspension by sample grinding with Dounce homogenizers (Sigma, catalog no. D8938-1SET) with 20 strokes with pestle A and 18 strokes with pestle B. The nuclei suspension was filtered by loading through a 35 μm diameter filter and followed by centrifuging 5 min at 600 g and 4°C. The nuclei pellet was collected and washed with cold wash buffer, which consisted of the following reagents: 1X PBS (Corning, catalog no. 46-013-CM), 20 mM DTT (Thermo Fisher Scientific, catalog no. P2325), 1%BSA (NEB, catalog no. B9000S), 0.2U/μl RNase inhibitor (Ambion, catalog no. AM2682) for three times. After removing the supernatant from the last wash, the nuclei were resuspended in 1 mL of 0.5 ug/ml DAPI (Sigma, catalog no. D9542) containing wash buffer to stain for 15 min. The nuclei suspension was prepared for sorting by filtering cell aggregates and particles out with a diameter of 35 μm. After removing myelin and fractured nuclei by sorting, the nuclei were collected by centrifuging 5 min at 600 g and 4°C, then resuspended in wash buffer to reach a final concentration of 1×10e6 nuclei/ml after counting in trypan blue (Thermo Fisher Scientific, catalog no. T10282) using a Countess II cell counter (Thermo Fisher Scientific, catalog no. A27977).

A single nuclei suspension containing 10,000 nuclei was loaded on a Chromium Single Cell B Chip (10x Genomics, catalog no. 1000154) as follows: 75 μl of master mix + nuclei suspension was loaded into the row labeled 1, 40μl of Chromium Single Cell 3’
s Gel Beads (10x Genomics, catalog no. PN-1000093) into the row labeled 2 and 280 μl of Partitioning Oil (10x Genomics, catalog no. 2000190) into the row labeled 3. This was followed by GEM generation and barcoding, post GEM-RT cleanup and cDNA Amplification. Then 100 ng purified cDNA derived from 12 cycles of cDNA amplification was used for 3’ Gene Expression Library Construction by using Chromium Single Cell 3’ GEM, Library & Gel Bead Kit v3 (10x Genomics, catalog no. 1000092) according to the manufacturer’s manual (10x Genomics, catalog no. CG000183 Rev C). The barcoded short-read libraries were measured using a Qubit 2.0 with a Qubit dsDNA HS assay kit (Invitrogen, catalog no. Q32854) and the quality of the libraries assessed on a Fragment analyzer (Agilent) using a high-sensitivity NGS Fragment Kit (1-6000bp) (Agilent, catalog no. DNF-474-0500). Sequencing libraries were loaded on an Illumina NovaSeq6000 with PE 2 × 50 paired-end kits by using the following read length: 28 cycles Read1, 8 cycles i7 index and 91 cycles Read2.

### Linear/asymmetric PCR steps to remove non-barcoded cDNA

The first round PCR protocol (95°C for 3 min, 12 cycles of 98°C for 20 s, 64°C for 30 s and 72°C for 60 s) was performed by applying 12 cycles of linear/asymmetric amplification to preferentially amplify one strand of the cDNA template (30 ng cDNA generated by using 10x Genomics Chromium Single Cell 3’
s GEM kit) with primer “Partial Read1”, then the product was purified with 0.8X SPRIselect beads (Beckman Coulter, catalog no. B23318) and washed twice with 80% ethanol. The second round PCR is performed by applying 4 cycles of exponential amplification under the same condition with forward primer “Partial Read1” and reverse primer “Partial TSO”, then the product was purified with 0.6X SPRIselect beads and washed twice with 80% ethanol and eluted in 30 ul buffer EB (Qiagen, catalog no. 19086). Sequences of primers: Partial Read1 (5’-CTACACGACGCTCTTCCGATCT-3’), Partial TSO (5’-AAGCAGTGGTATCAACGCAGAGTACAT-3’). KAPA HiFi HotStart PCR Ready Mix (2X) (Roche, catalog no. KK2601) was used as polymerase for all the PCR amplification steps in this paper except for the single nuclei RNA-Seq cDNA amplification and library construction part.

### Exome capture to enrich for spliced cDNA

For each sample, exome enrichment was applied to the cDNA purified from the previous step to enrich spliced cDNA by using probe kit SSELXT Human All Exon V8 (Agilent, catalog no. 5191-6879) and the compatible reagent kit SureSelectXT HSQ (Agilent, catalog no. G9611A) according to the manufacturer’s manual. Firstly, the block oligo mix was made by mixing equal amount (1 μl of each per reaction) of primers Partial Read1 (5’-CTACACGACGCTCTTCCGATCT-3’) and Partial TSO (5’-AAGCAGTGGTATCAACGCAGAGTACAT-3’) with the concentration of 200 ng/μl (IDT), resulting in a final concentration of 100 ng/μl. Next, 5 μl of 100 ng/μl cDNA diluted from the previous step was combined with 2 μl block mix and 2 μl nuclease free water (NEB, catalog no. AM9937), then the cDNA-block oligo mix was incubated on a thermocycler under the following condition to allow block oligo mix to bind to 5’ and 3’ end of the cDNA molecule: 95°C for 5 min, 65°C for 5 min, 65°C on hold. For the next step, the hybridization mix was prepared by combining 20 ml SureSelect Hyb1, 0.8 ml SureSelect Hyb2, 8.0 ml SureSelect Hyb3, and 10.4 ml SureSelect Hyb4 and kept at room temperature. Once the reaction reached to 65°C on hold, 5 μl of probe (SSELXT Human All Exon V8), 1.5 μl of Nuclease free water, 0.5 μl of 1:4 diluted RNase Block (SureSelectXT HSQ) and 13 μl of the hybridization mix were added to the cDNA-block oligo mix and incubated for 24 hrs at 65°C. When the incubation reached the end, the hybridization reaction was transferred to room temperature. At the same time, an aliquot of 75 μl M-270 Streptavidin Dynabeads (Thermo Fisher Scientific, catalog no. 65305) were prepared by washing three times with binding buffer (SureSelectXT HSQ) and resuspended in 200 μl binding buffer. Next, the hybridization reaction was mixed with all the M270 Dynabeads and placed on a Hula mixer with a high-speed for 30 mins at room temperature. During the incubation, 600 μl of wash buffer 2 (SureSelectXT HSQ) was transferred to 3 wells of 0.2 ml 8-strip tube and incubated in a thermocycler on hold at 65°C. When the 30 min incubation was over, the buffer was replaced with 200 μl of wash buffer 1 (SureSelectXT HSQ). Then the tube containing hybridization product bound to M-270 Dynabeads was put back to the Hula mixer for another 15 min incubation with low speed. Next, the wash buffer 1 was replaced with wash buffer2 and the tube was transferred to the thermocycler for the next round of incubation. Overall, the hybridization product bound to M-270 Dynabeads was incubated in wash buffer 2 for 30 min at 65°C, and the fresh pre-heated wash buffer 2 was applied every 10 min. For the last step, the wash buffer 2 was removed when the incubation was over and the beads were resuspended in 18 μl of Low TE buffer and stored in 4°C. Next, the spliced cDNA which bound with the M-270 Dynabeads was amplified with primers Partial Read1 and Partial TSO by using the following PCR protocol: 95°C for 3 min, 12 cycles of 98°C for 20 s, 64°C for 60 s and 72°C for 3 min. Then the amplified spliced cDNA was isolated from M-270 beads as supernatant, and followed by a purification with 0.6✕ SPRIselect beads and washed twice with 80% ethanol.

### Library preparation for PacBio

HiFi SMRTbell Libraries of FCtx1 and FCtx2 were constructed according to the manufacturer’s manual (entitled “Preparing HiFi SMRTbell Libraries using SMRTbell Express Template Prep Kit 2.0”) by using SMRTbell Express Template Prep Kit 2.0 (PacBio, catalog no. 100-938-900). For both samples, ∼500 ng cDNA obtained by performing LAP-CAP and Exome enrichment from the previous step was used for library preparation. The library construction includes: DNA damage repair (37°C for 30 mins), end-repair/A-tailing (20°C for 30 mins, 65°C for 30 mins), adaptor ligation (20°C for 60 mins), and purification with 0.6X SPRIselect beads and washed twice with 80% ethanol.

### Library preparation for ONT

For both samples, approximately 50-75 fmol cDNA processed through LAP-CAP and exome enrichment was used for ONT library construction by using Ligation Sequencing Kit (Oxford Nanopore Technologies, catalog no. SQK-LSK110) according to the manufacturer’s protocol (Nanopore Protocol, Amplicons by Ligation, Version: ACDE_9110_v110_revC_10Nov2020). First, cDNA was subjected to DNA repair and end-prep using the NEBNext FFPE DNA Repair Mix and NEBNext Ultra II End repair / dA-tailing Module reagents (NEB, catalog no. M6630 and E7546) in accordance with the manufacturer’s instructions. After incubating at 20°C for 5 mins and 65°C for 5 mins, cDNA was purified with 1X SPRIselect beads and washed twice with 70% ethanol. Then, ONT adapters were ligated using the NEBNext T4 Quick Ligase (NEB, catalog no. NEBNext Quick Ligation Module, E6056) and incubated for 10 min at room temperature, followed by purification with 0.4X SPRIselect beads and washing with SFB buffer. The ONT library from the last step was loaded onto a PromethION sequencer by using PromethION Flow Cell (Oxford Nanopore Technologies, catalog no. FLO-PRO002) and sequenced for 72 hrs. Base-calling was performed with Guppy by setting base-quality score >7.

### Data processing and QC for single-cell short-read analysis

The 10x cellranger pipeline (version 3.1.0) was run on the raw Illumina sequencing data to obtain single-cell expression matrices which were then analyzed using Seurat v3.1.1^31^. For both FCtx samples, nuclei that had unique gene counts over 7,500 or less than 200, and greater than 4% mitochondrial gene expression were removed from further analysis. Filtering on these parameters yielded 7,314 single nuclei for the FCtx1 sample, and 6,486 single nuclei for FCtx2. The number of UMIs and percentage of mitochondrial gene expression were regressed from each nucleus and then the gene expression matrix was log normalized and scaled to 10,000 reads per cell. Next, we clustered cells using the Louvain algorithm, setting the resolution parameter to 0.6. To visualize and explore these datasets, we performed both tSNE and UMAP non-linear reduction techniques. Cell types were assigned by identifying canonical marker genes for each cluster^13,45–47^. This cell type annotation was confirmed by aligning to the Allen Brain Atlas human cortical data^13^.

### Alignment of PacBio long-read data

Using the default SMRT-Link parameters, we performed circular consensus sequencing (CCS) with IsoSeq3 with the following modified parameters: maximum subread length 14,000 bp, minimum subread length 10 bp, and minimum number of passes 3.

Long read CCS fastq sequences with PacBio were mapped and aligned to the reference genome (GRCh38) using STARlong and the following parameters:

--readFilesCommand zcat --runMode alignReads --outSAMattributes NH HI NM MD -- readNameSeparator space --outFilterMultimapScoreRange 1 --outFilterMismatchNmax 2000 -- scoreGapNoncan -20 --scoreGapGCAG -4 --scoreGapATAC -8 --scoreDelOpen -1 --scoreDelBase -1 -- scoreInsOpen -1 --scoreInsBase -1 --alignEndsType Local --seedSearchStartLmax 50 -- seedPerReadNmax 100000 --seedPerWindowNmax 1000 --alignTranscriptsPerReadNmax 100000 - -alignTranscriptsPerWindowNmax 10000.

### Alignment of ONT long-read data

Long reads sequenced on the ONT PromethION were mapped and aligned using minimap2 (v2.17- r943-dirty) using the following parameters: -t 20 -ax splice --secondary=no.

### Barcode detection and identification of unique molecules from PacBio data

Cellular barcodes (16 nt) were obtained for each single-nucleus assigned to a cell type. Assuming an error rate of 10%, PolyA tails were identified by obtaining the position of 9 consecutive A’s or T’s in the last 100 nt of PacBio CCS and ONT reads respectively. Leveraging the position of the PolyA tail, a sliding window was used to identify perfect matching barcodes in the 36 bp window preceding the PolyA tail, and UMIs were identified as the 12 bp sequence immediately following the barcode. This method is implemented in the ‘scisorseqr’ package under the ‘GetBarcodes’ function. Given some amount of PCR duplication, one transcript per molecule i.e. barcode+UMI+gene was chosen for analysis.

### Barcode detection for long read transcripts obtained from ONT

Perfect matching barcodes were obtained in a similar fashion to the PacBio reads described above. However, given the higher error rate of ONT data, we wanted to allow for more molecules with sequencing errors. Therefore, using the mapping information per read along with a white-list of UMIs as done before by others^48^, we modified our strategy as follows:

- For each molecule sequenced on the 10x, we obtained a 28 bp barcode-UMI sequence, and grouped them by gene (Supplementary Fig. 8a).
- This was a comprehensive set of reference molecules we would use to identify barcode-UMI sequences from the ONT long reads.
- For every ONT read mapped to a gene, we narrowed the search space by using only the reference list for that gene (Supplementary Fig. 8b).
- We used a sliding window approach to identify potential barcode-UMI sequence in the long reads by allowing for up to 1 mismatch in the first 22 bp of each molecule in the reference list. We then filtered the potential matches by allowing for either 1 or 2 mismatches in the 28 bp identified sequences (Supplementary Fig. 8c-d).
- The 1-2 mismatches were allowed at this step because the collision rate of barcode sequences from the 10x output was low, and 10x UMIs showed an uncharacteristically higher collision rate for an edit distance of 2 and we did not want to discard molecules that had a high likelihood of being legitimate. (Supplementary Fig. 8e-f).

Therefore, we allowed for up to two mismatches in the barcode-UMI pairs from the list of molecules obtained from the short-read analysis. These steps were performed using a custom script.

### Identification of unique molecules from ONT data

Given the higher error rate of ONT sequencing output, the reads were more likely to undergo “molecule inflation”, meaning that a single error in a read could result in one molecule being perceived as two different ones and therefore inflate our counts (Supplementary Fig. 9a). To combat this issue, we corrected for sequencing errors in reads. This was performed as follows:

- Reads were grouped by barcode-UMI-gene and ordered by frequency with the highest frequency indicating PCR duplicates of the same molecule.
- The edit distance to the nearest barcode-UMI-gene pair from the short-read reference set was obtained using the Levenshtein distance.
- If the distance to the reference was zero, i.e. it was a perfect match, the molecule was retained.
- If the distance was either 1 or 2, the 28 bp sequence was corrected to match the reference, and if the molecule was novel with respect to ONT data, then it was retained (Supplementary Fig. 9b, left).
- If the distance to the reference was greater than 3, and the edit distance to any other already accepted UMI was sufficiently large, i.e. greater than 5, the UMI was deemed to be novel and was retained (Supplementary Fig. 9b, purple box).
- If the distance to the reference was greater than 3, but the edit distance to any other already accepted UMI was either 1 or 2, then it was deemed to have a sequencing error and the UMI was corrected to the accepted UMI (Supplementary Fig. 9b, red box, right).

Following the sequencing error correction according to these above metrics, only one transcript per molecule i.e. barcode + UMI + gene was chosen for further analysis. These steps were performed using a custom script.

### Assignment of TSS and PAS to reads

TSS and PAS were assigned to each read as previously done. Specifically, we assigned the closest previously published TSS within 50 bp of the 5’ end of the read mapping^49^. Likewise, we assigned the closest published PAS within 50 bp of the 3’ end of the read mapping^50^.

### Obtaining phastCons scores for exons, TSS, PolyA sites from 17 primates

PhastCons scores for 16 primate genomes aligned to the human genome were obtained from the UCSC website^41,51^. The scores were then averaged over specified genomic regions (i.e. internal exons, TSS, and PAS that were identified as described above) using the *bigWigAverageOverBed* script from the UCSC Utilities package.

### Obtaining lists of disease associated exons for ASD, schizophrenia, ALS

The list of ASD associated exons which covers 3,482 unique exons were summarized from two genome-wide studies and one review: 1,776 skipped exons detected in analysis of differential splicing changes in human cortex by comparing ASD cases with controls (p-value<0.05)^36^, 1,723 neural regulated alternative spliced exons detected in autistic human brains^34^, and 33 unique microexons which are associated with ASD and also characterized at the functional leve^l37^ were collected. For Schizophrenia, a list of Schizophrenia associated exons which were classified as alternative exon skipping events covering 1,107 exons were collected^35^. The list of 506 ALS associated cassette exons identified by comparing *C9orf72* ALS brains with control brains were collected^38^.

### Alternative exon counting and categorization

Starting with all exons that appeared as an internal exon in a read, we calculated the following numbers:

1. The number of long-read UMIs containing this exon with identity of both splice sites: *X*_*in*_
2. The number of long-read UMIs assigned to the same gene as the exon, which skipped the exon as well as 50 adjacent bases on both sides: *X*_*out*_
3. The number of long-read UMIs supporting the acceptor of the exon and ending on the exon itself: *X*_*acc*_*In*_
4. The number of long-read UMIs supporting the donor of the exon and ending on the exon itself: *X*_*don*_*In*_
5. The number of long-read UMIs overlapping the exon: *X*_*tot*_

Non-annotated exons with one or two annotated splice sites, ≥70 bases of non-exonic (in comparison to the annotation) bases, were then excluded as intron-retention events or alternative acceptors or donors.

We then calculated

- 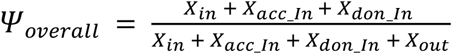
- 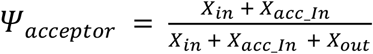
- 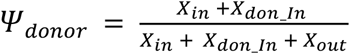

If

- 0.05 ≤ *Ψ*_*condition*_ ≤ 0.95 *where condition* ∈ {*overall, acceptor, donor*}
- 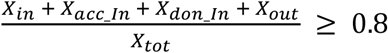

then the exon was kept for further analysis.

We then calculated the *Ψ*_*overall*_ for each cell type as 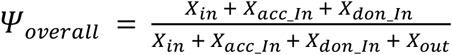 from all the long-read UMIs for that cell type if and only if *X*_*tot*_ ≥ 10 for the exon and cell type in question. When fewer reads were available the *Ψ*_*overall*_ for the exon and cell type in question were set to “NA”.

### Exon variability analysis

For each replicate, we defined a set of alternative exons that met each of the following criteria: (i) ≥10 supporting reads (inclusion + exclusion) at the pseudo-bulk level; (ii) 0.05 < *Ψ* < 0.95 at the pseudo-bulk level; and (iii) intron retention events were excluded. These filtering steps produced a set of 5,855 (FCtx1) and 5,273 (FCtx2) alternative exons. From this set, we further defined a subset of alternative exons that had ≥10 supporting reads in each of the four major cell-types (excitatory neurons, inhibitory neurons, astrocytes and oligodendrocytes). To investigate the relationship between exon length and inclusion variability amongst cell-types, we divided alternative exons into three variability categories: (i) (max*Ψ* - min*Ψ*) ≤ 0.25; (ii) 0.25 < (max*Ψ* - min*Ψ*) ≤ 0.75; and (iii) (max*Ψ* - min*Ψ*) > 0.75. For each category, we plotted the density of exon length using ggplot2. Next, we investigated whether exons associated with three neurological diseases (schizophrenia, ALS and ASD; see above) show different inclusion variability amongst cell types. For each disease, we compared disease-associated exons with all other alternative exons in our subset and performed a two-sided Wilcoxon rank sum test. Micro-exons were defined as exons with a length of ≤27 bp. Novel exons were defined as exons that are not described in the GENCODE v34 annotation. To define a subset of novel exons that show high inclusion and/or high cell-type variability, we plotted (max*Ψ* - min*Ψ*) against pseudo-bulk *Ψ* and fit a loess curve to the data.

### Testing for exon coordination

Testing for exon coordination can be done either by aggregating counts for all cell types and testing for association at the pseudo-bulk level, or at the individual cell type level. For every exon pair passing the aforementioned criteria for sufficient depth, a *2 × 2* matrix of association for a given sample i.e. cell type or pseudo-bulk was generated. This matrix contained counts for inclusion of both exons (in-in), inclusion of the first exon and exclusion of the second (in-out), exclusion of the first exon and inclusion of the second (out-in), and exclusion of both exons (out-out).

The co-inclusion score of an exon was defined as the double inclusion (in-in) divided by the total counts for that exon pair. An exon pair was deemed to be coordinated if counts for double inclusion or double exclusion were significantly higher than counts for mutually exclusive exons. Significance levels were obtained by calculating a p-value from the **χ**^2^ test of association. The effect size was calculated as the |log_10_(oddsRatio)| (LOR). The odds-ratio was calculated explicitly by setting 0 values to 0.5, and dividing the product of double inclusion and double exclusion by the product of single-inclusion i.e. (in-in) * (out-out) / (in-out) * (out-in). Finally, given that there are multiple exons per gene and therefore the tests for associations are dependent, we used a Benjamini-Yekutieli correction for multiple testing and reported the FDR value.

### Conservation analysis for exon pairs

PhastCons scores from primates were obtained as mentioned above. For every gene used in the pseudo-bulk analysis, the exon pairs with the smallest |log_10_(oddsRatio)| were chosen for further analysis. Once these exon pairs were defined, the minimum phastCons score for each pair was extracted and plotted against the absolute value of the LOR.

### Cell-type specific conservation analysis for exon pairs

Exon coordination count data was then split by cell type, including astrocyte, excitatory neuron, inhibitory neuron, oligodendrocyte, or oligodendrocyte precursor cells (OPCs). In conjunction with the LOR, here we calculated an exon inclusion ratio which is defined as the number of times both exons pairs are included in the sequencing data (in-in), divided by the coordination i.e. sum of the in-in and out-out counts per exon pair. The minimum phastCons value for each exon pair was selected and placed into 1 of 2 groups based on value. The first group included all phastCons values below 0.5, and the second contained values above 0.5. We then plotted these two groups against the |log_10_(oddsRatio)| as well as the exon inclusion ratio as box plots.

### Obtaining counts for exon – end site combinations

We obtained counts for exon-TSS and exon-PAS combinations using a custom script. Specifically, if a read was assigned to a TSS and contained an internal exon, then it was counted as including the exon for that TSS. On the other hand, if it was assigned to a TSS and the read skipped the internal exon even though the read was long enough to cover it, then it was counted as excluding the exon for the TSS. The same principle was applied for counting inclusion and exclusion levels of exon-PAS combinations. Only genes with sufficient depth i.e. at least 25 reads per gene were used in the count generation and then used for further analysis.

### Testing for exon – end site coordination

Similar to testing for exon coordination, testing for exon – end site coordination can be done either by aggregating counts for all cell types and testing for association at the pseudo-bulk level, or at the individual cell type level. For each such sample, a matrix of association was generated. This matrix was an *n x 2* table per internal exon, where the *n* TSS associated with the exon formed rows, while inclusion and exclusion counts formed columns. A p-value of differential usage was obtained from a **χ**^2^ test. For effect size, we used the metric of ***ΔΠ***, which is the maximum change in TSS inclusion levels. This was done by first calculating the percent inclusion (PI) and percent exclusion (PE) for each TSS as a percentage of total inclusion and exclusion counts respectively. The TSS with the maximum difference in PI and PE was therefore the most differentially used TSS, with the difference being reported as the ***ΔΠ***. The same approach was used for the PAS-exon coordination analysis. Finally, given that there are multiple exons per gene and therefore the tests for associations are dependent, we used a Benjamini-Yekutieli correction for multiple testing and reported the FDR value.

### Definition of the χ^2^ criterion and classifying untestability due to constitutiveness

To categorize exon pairs or exon – end site tables as being untestable due to being constitutive, we employed the following metrics. For each matrix M with elements m_ij_ where m is in the i^th^ row and j^th^ column,

- The expected value for each element m_ij_ was defined as 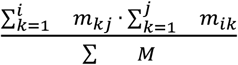.
- If the expected value in 80% (rounded to nearest integer) of elements is at least 5, and the expected value of all element is at least 1, then the **χ**^2^ criterion is met and the p-value is calculated
- If the median expected value is < 5 in any row or any column, then the RNA-variable (i.e. TSS, exon, or PAS) in that row or column is said to be constitutive.

### Conservation analysis for exon – end site pairs

PhastCons scores for all defined TSS were extracted as described above. For every gene used in the pseudo-bulk analysis, the TSS-exon pair with the smallest **Δ*Π*** was chosen. For each of these exons, phastCons scores of the associated TSS were sorted by value. The TSS with the minimum phastCons score was reported for that exon, and the Pearson’s product-moment correlation between the phastCons score and **Δ*Π*** for that TSS-exon pair was calculated. Similar analysis was conducted for the exon-PAS pairs.

## Data availability

All data used for this study will be made publicly available on GEO under the accession token GSE178175. All data supporting the findings of this study are provided within the paper and its supplementary information. All additional information will be made available upon reasonable request to the authors.

## Code availability

The source code generated for this paper will be made publicly available at https://github.com/noush-joglekar/sn-code.

## Supplementary figure legends

**Supplementary Figure 1: Cell type assignment for frontal cortex single-nuclei sample. a**. UMAP plot of FCtx1 sample with clusters labelled by cell type annotation. **b**. Heatmap of gene expression of marker genes that are enriched in each cluster depicted in A, with purple denoting low expression while yellow denotes high expression. Each row corresponds to a gene while each column corresponds to a single nucleus. Nuclei are clustered by cell type, and annotation for each cell type is at the top. **c-r**. Normalized gene expression for the indicated marker genes projected onto the UMAP plot.

**Supplementary Figure 2: Short-read sequencing statistics. a-b**. Violin plots depicting UMIs (a) and genes (b) sequenced per single nucleus broken down by cell type which is indicated on the X-axis. **c**. Barplot of the percentage of total sequenced reads assigned to each of the metrics defined on the X-axis. Color of bar indicates the sample. **d**. Barplot of the number of single nuclei assigned to each cell type indicated on the X-axis. Color of bar indicates the sample.

**Supplementary Figure 3: Long-read sequencing statistics. a-b**. Histogram of reads per single nucleus with reads on the X-axis and number of single nuclei sequenced on the Y-axis. FCtx1 on the left (red) and FCtx2 on the right (blue). **c-d**. Barplot of the number of single-nuclei recovered per cell type. Color of bar indicates sample, i.e. FCtx1 on the left and FCtx2 on the right. **e-f**. Boxplots of reads per single nucleus, grouped by cell type. **g-h**. Boxplots of UMIs per single nucleus, grouped by cell type. **i-j**. Boxplots of genes per single nucleus, grouped by cell type, with FCtx1 on the left and FCtx2 on the right.

**Supplementary Figure 4: Replicable observations for single exon usage. a-d**. Panels correspond to Fig. 3b-e, but using data from FCtx2.

**Supplementary Figure 5: Replicable coordination of adjacent and distant exon pairs. a-b**. Barplot showing percent of tested genes in pseudo-bulk with significant exon coordination (FDR≤0.05) for adjacent (a) and distant (b) exon pairs at various log-odds ratio (LOR) cutoffs on the X-axis. Error bars indicate standard error of the point estimate. **c**. Boxplots of the absolute value of LOR for significant genes (FDR ≤ 0.05) on the Y-axis plotted against adjacent and distant exon pairs (A-B) on the X-axis. p-value obtained from two-sided Wilcoxon rank sum test. **d**. Density plot for the LOR for adjacent and distant exon pairs. **e**. Barplots showing percent of tested genes (FDR≤0.05, abs(LOR) ≥1) in pseudo-bulk for distant exons pairs on the X-axis where either one, or both, or neither of the exons are known to be associated with ASD. Error bars indicate standard error of the point estimate and p-values obtained from two-sided Fisher’s exact test.

**Supplementary Figure 6: Exon coordination patterns across multiple cell types are replicable**.

**a**. Pie chart indicating the number of cell types where an exon pair is significant (FDR ≤ 0.05 and abs(LOR)≥1) given that the exon-pair is testable in at least one cell type, and is significant when tested in pseudo-bulk. **b**. Barplot indicating percentage of tested exon-pairs (one per gene) that were deemed significantly coordinated (FDR ≤ 0.05 and abs(LOR) ≥ 1). Breakdown by cell type on the X-axis, given significance in pseudo-bulk. Error bars represent the standard error of the point estimate. **c**. Heatmap showing cell types as columns and exon pairs that were testable in at least one cell type as rows. Each element of the heatmap is colored by whether the exon-pair showed significant coordination and the direction of coordination with respect to the pseudo-bulk (same: pink or opposite: blue), was not significant (white), was not testable in a particular cell type because it had too few counts to satisfy the **χ**^2^ criterion (grey) or because one or both exons became constitutively included in a particular cell type (teal). **d**. Boxplots of the absolute value of the LOR for tested exon pairs for the excitatory neuron reads denoted on the Y-axis and split by whether the minimum phastCons score of the two exons was greater than or less than 0.5. Significance calculated using two-sided Wilcoxon rank sum test. **e**. Boxplots for the excitatory neuron reads showing the double inclusion level of an exon pair divided by the number of reads that were coordinated i.e. double inclusion + double exclusion. X-axis indicated whether the minimum phastCons score of the two exons was greater than or less than 0.5. Significance was calculated using the two-sided Wilcoxon rank sum test.

**Supplementary Figure 7: Cell type mediated exon - end site coordination in FCtx2. a**. Bar chart indicating the number of cell types where an exon-end site pair is significant (FDR ≤ 0.05 and **ΔΠ**≥1) given that the exon-end site pair is significant when tested in pseudo-bulk and is testable in at least one cell type. Color of bar indicates whether the end site associated with an exon is a TSS (blue) or PAS (green). Error bars represent standard error of the point estimate. **b**. Bar chart showing percent of tested genes in pseudo-bulk with significant exon-end site coordination (FDR ≤ 0.05) on the Y-axis and various ***ΔΠ*** cutoffs on the X-axis. Color of bar indicates whether the end site associated with an exon is a TSS (blue) or PAS (green). Error bars indicate standard error of the point estimate. **c-d**. Heatmaps showing cell types as columns and exon-end site pairs (exon-TSS left, exon-PAS right) that were testable in at least one cell type as rows. Each element of the heatmap is colored by whether the exon-end site pair showed significant coordination (pink), was not significant (white), was not testable in a particular cell type because it had too few counts to satisfy the **χ**^2^ criterion (grey), or because the exon or one or more end sites became constitutively included in a particular cell type (teal).

**Supplementary Figure 8: Barcode detection for long read transcripts obtained from ONT. a**.

Reference set of molecules (Barcode-UMI sequence) for *GeneX* from 10x short-read sequencing data. **b**. ONT reads mapped to reference *GeneX*. **c**. First 22 bp of 28 bp long reference sequence used as a search space. **d**. Isolating ONT reads with up to 1 mismatch in 22 bp of reference sequence shown in C. Getting the full 28 bp barcode + UMI sequence and only retaining reads with up to 2 mismatches. **e**. Histogram of Levenshtein distances of every pairwise comparison of barcodes found in 10x FCtx1 sample. **f**. Histogram of Levenshtein distances of every pairwise comparison of UMIs per gene-barcode pair reported in 10x FCtx1 sample. Red box highlights a higher than expected rate for 2 mismatches within UMIs for a single gene-barcode pair.

**Supplementary Figure 9: Identification of unique molecules from ONT data. a**. Illustration of ‘molecule inflation’ caused by PCR duplication and sequencing errors in ONT data for a single barcode-UMI pair associated with a gene. Histogram of Levenshtein distances of every pairwise comparisons of UMIs per gene-barcode pair found in the FCtx1 sample. *Left:* comparisons of ONT UMIs with the 10x reference set. *Right:* comparisons of ONT UMIs with other ONT UMIs associated with a gene-barcode pair. Red boxes highlight the fraction of UMIs that were corrected to the reference. Purple box highlights the UMIs which were sufficiently different from all existing UMIs and were therefore considered to be novel.

