## Supplementary Figures 1-9 and Supplementary Table 1 for "Single-nuclei isoform RNA sequencing reveals combination patterns of transcript elements across human brain cell types"

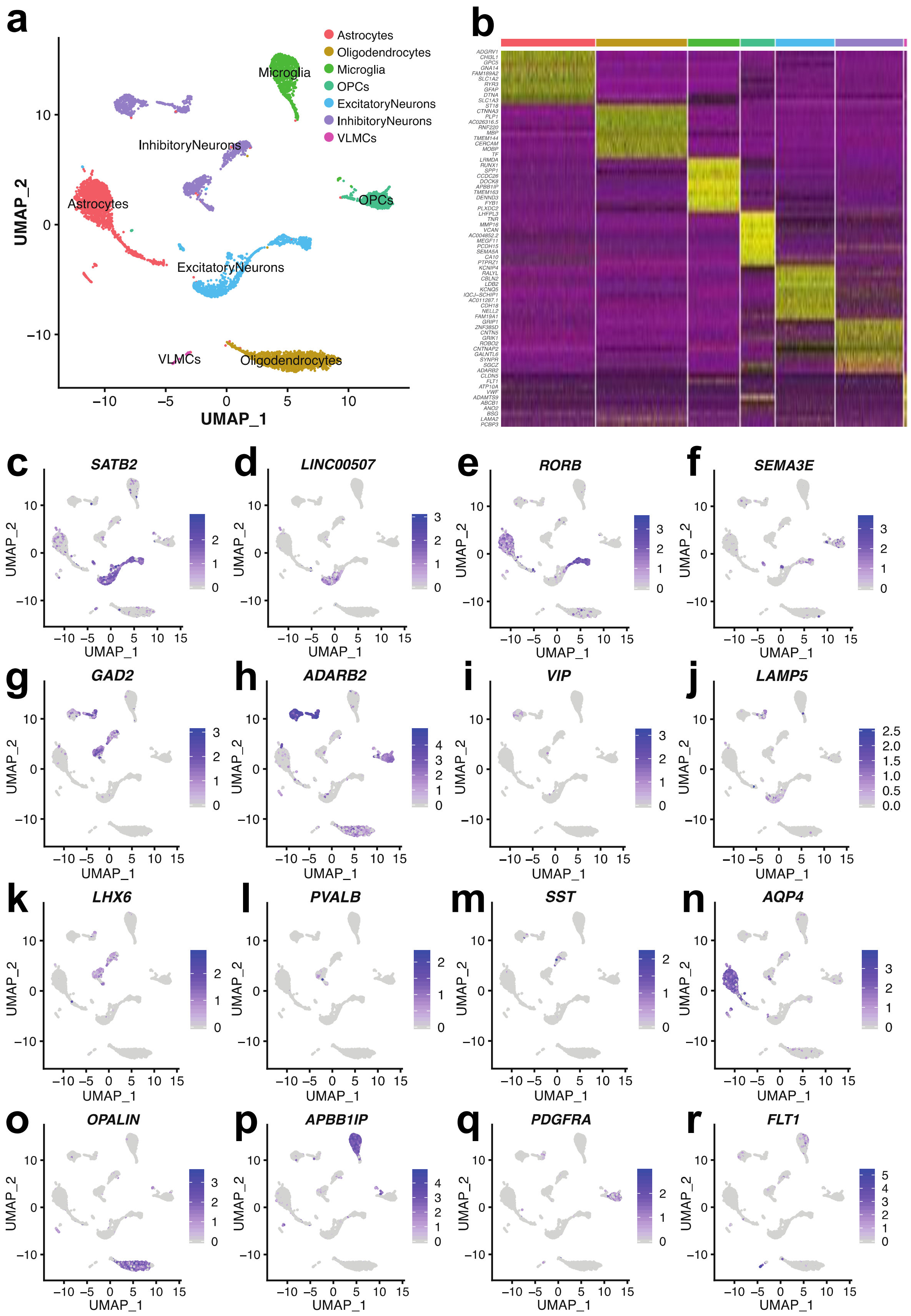


**Supplementary Figure 1: Cell type assignment for frontal cortex single-nuclei sample. a.** UMAP plot of FCtx1 sample with clusters labelled by cell type annotation. **b.** Heatmap of gene expression of marker genes that are enriched in each cluster depicted in A, with purple denoting low expression while yellow denotes high expression. Each row corresponds to a gene while each column corresponds to a single nucleus. Nuclei are clustered by cell type, and annotation for each cell type is at the top. **c-r.** Normalized gene expression for the indicated marker genes projected onto the UMAP plot.

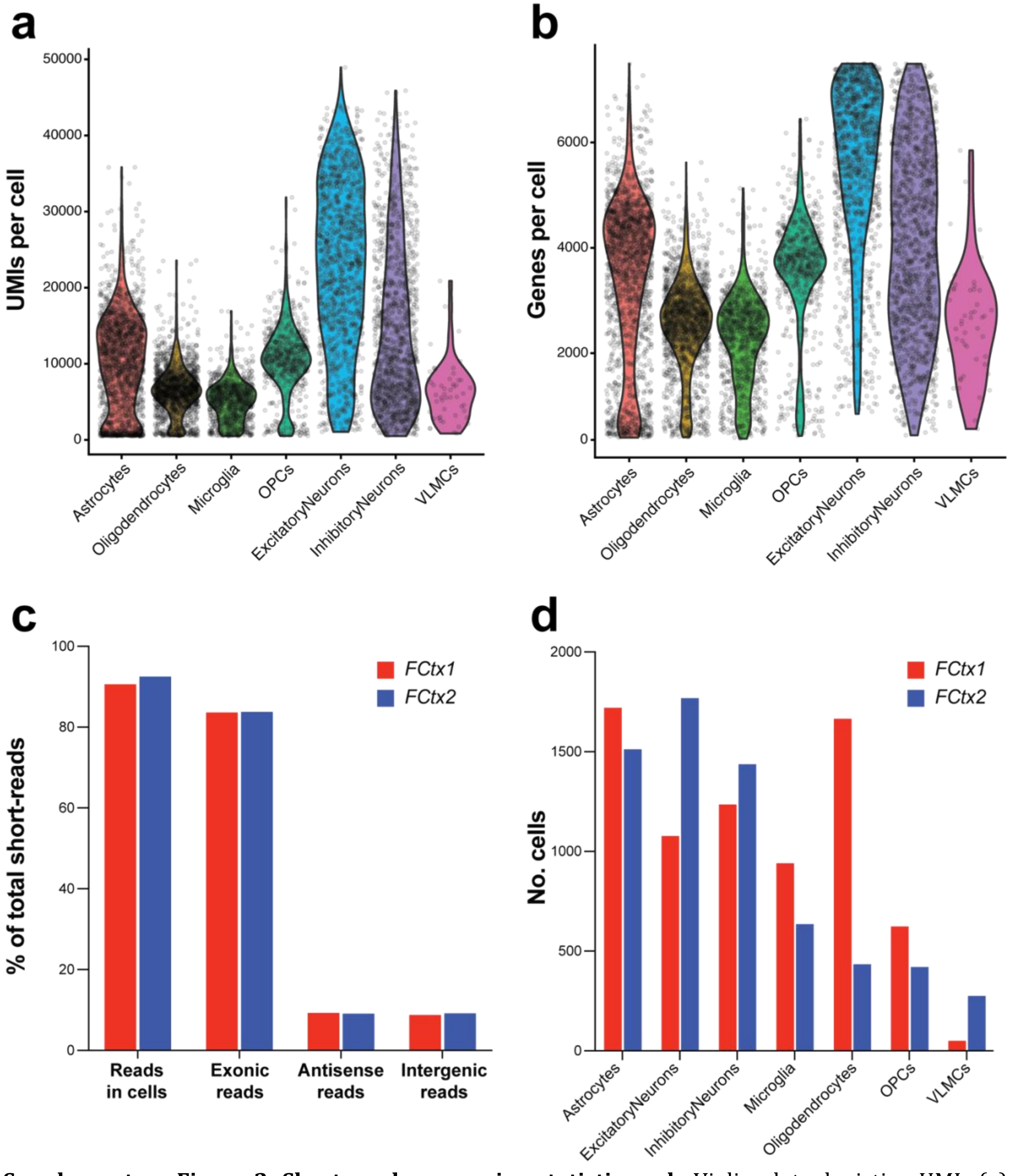


**Supplementary Figure 2: Short-read sequencing statistics. a-b.** Violin plots depicting UMIs (a) and genes (b) sequenced per single nucleus broken down by cell type which is indicated on the X-axis. **c.** Barplot of the percentage of total sequenced reads assigned to each of the metrics defined on the X-axis. Color of bar indicates the sample. **d.** Barplot of the number of single nuclei assigned to each cell type indicated on the X-axis. Color of bar indicates the sample.

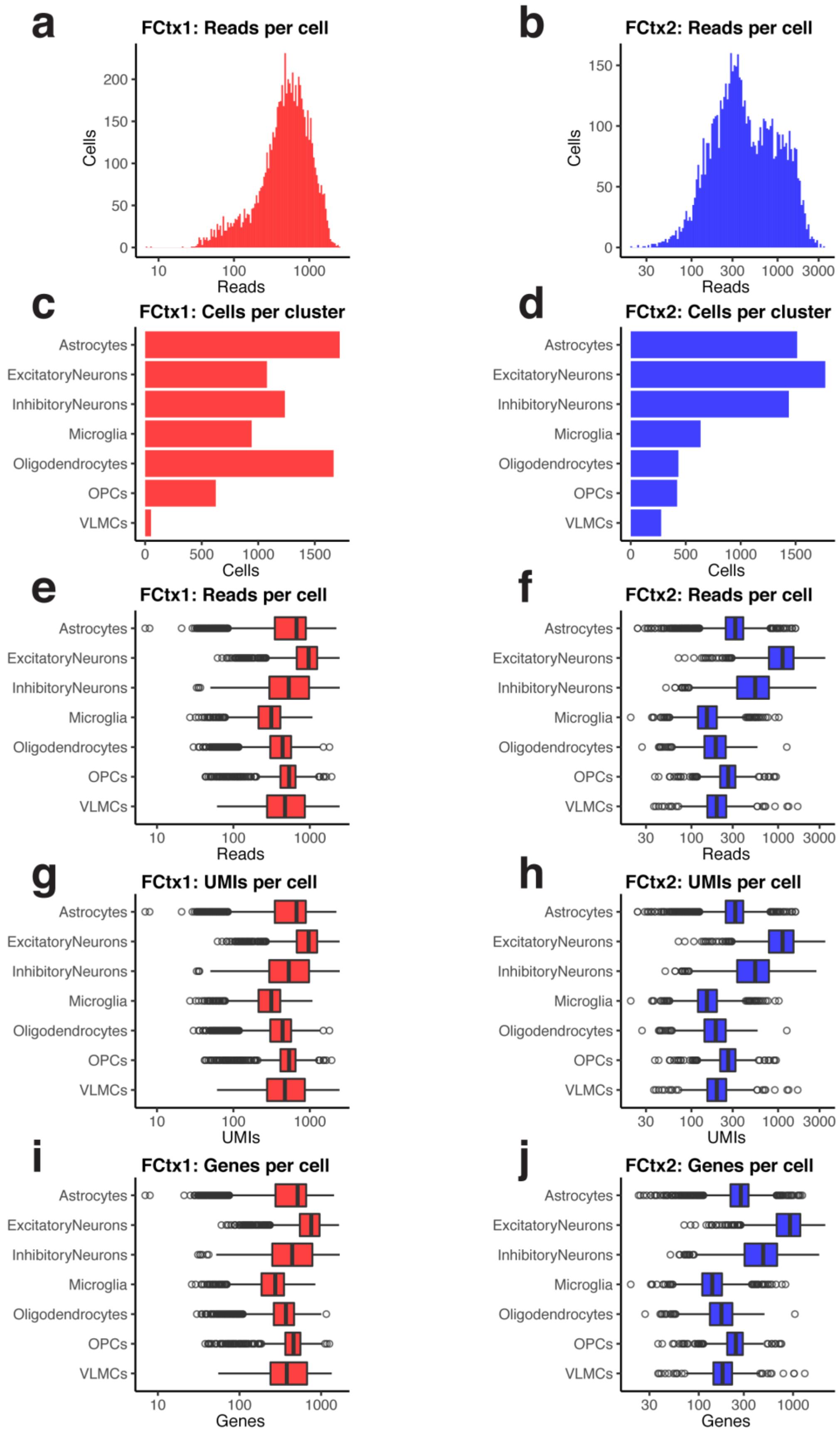
a
FCtx1: Reads per cell
Cells
Reads
b
FCtx2: Reads per cell
Cells
Reads
c
FCtx1: Cells per cluster
d
FCtx2: Cells per cluster
Astrocytes
ExcitatoryNeurons
InhibitoryNeurons
Microglia
Oligodendrocytes
OPCs
VLMCs
Cells
e
FCtx1: Reads per cell
f
FCtx2: Reads per cell
Reads
g
FCtx1: UMIs per cell
h
FCtx2: UMIs per cell
UMIs
i
FCtx1: Genes per cell
j
FCtx2: Genes per cell
Genes

**Supplementary Figure 3: Long-read sequencing statistics. a-b.** Histogram of reads per single nucleus with reads on the X-axis and number of single nuclei sequenced on the Y-axis. FCtx1 on the left (red) and FCtx2 on the right (blue). **c-d.** Barplot of the number of single-nuclei recovered per cell type. Color of bar indicates sample, i.e. FCtx1 on the left and FCtx2 on the right. **e-f.** Boxplots of reads per single nucleus, grouped by cell type. **g-h.** Boxplots of UMIs per single nucleus, grouped by cell type. **i-j.** Boxplots of genes per single nucleus, grouped by cell type, with FCtx1 on the left and FCtx2 on the right.

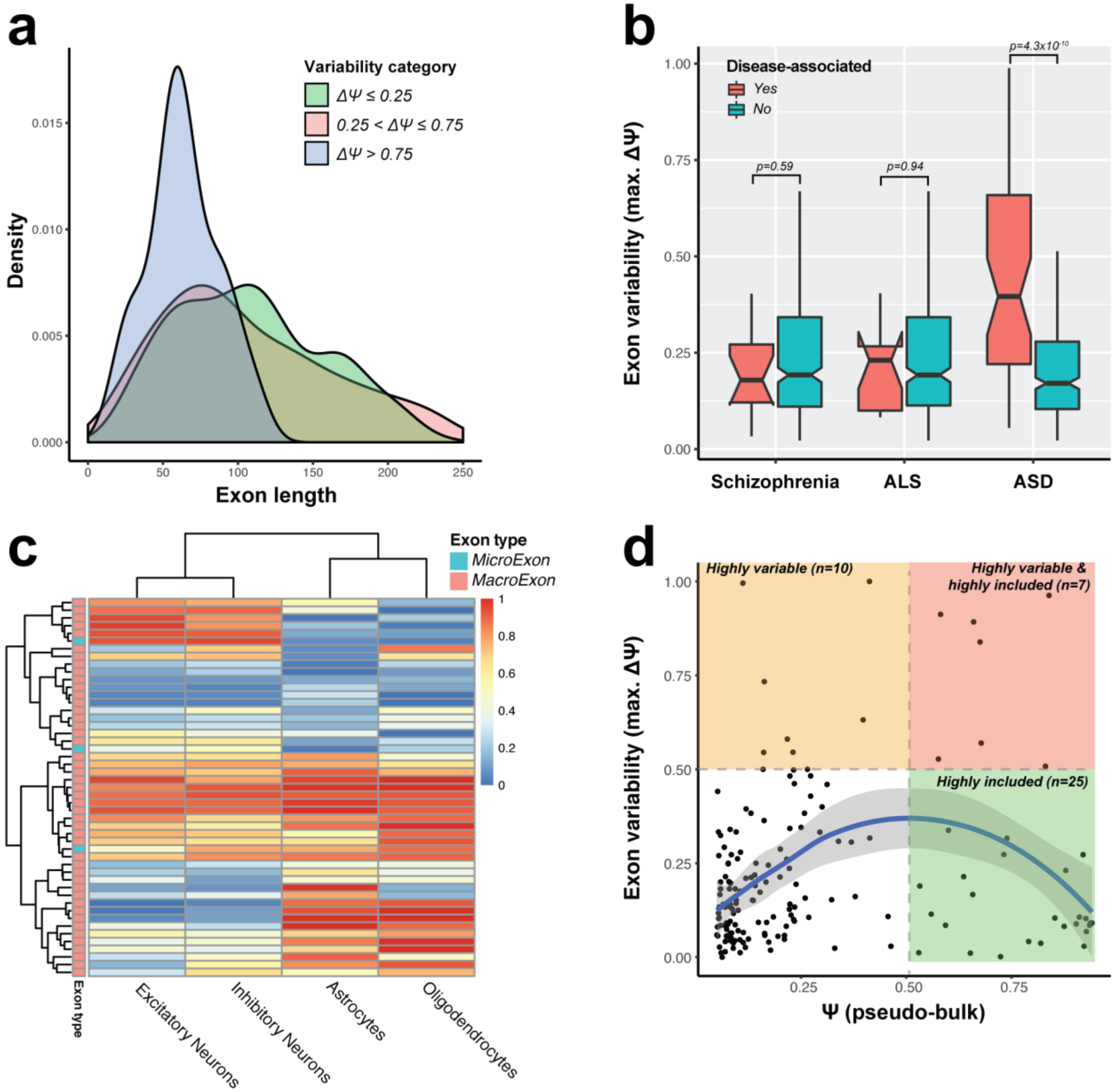


**Supplementary Figure 4: Replicable observations for single exon usage. a-d.** Panels correspond to Fig. 3b-e, but using data from FCtx2.

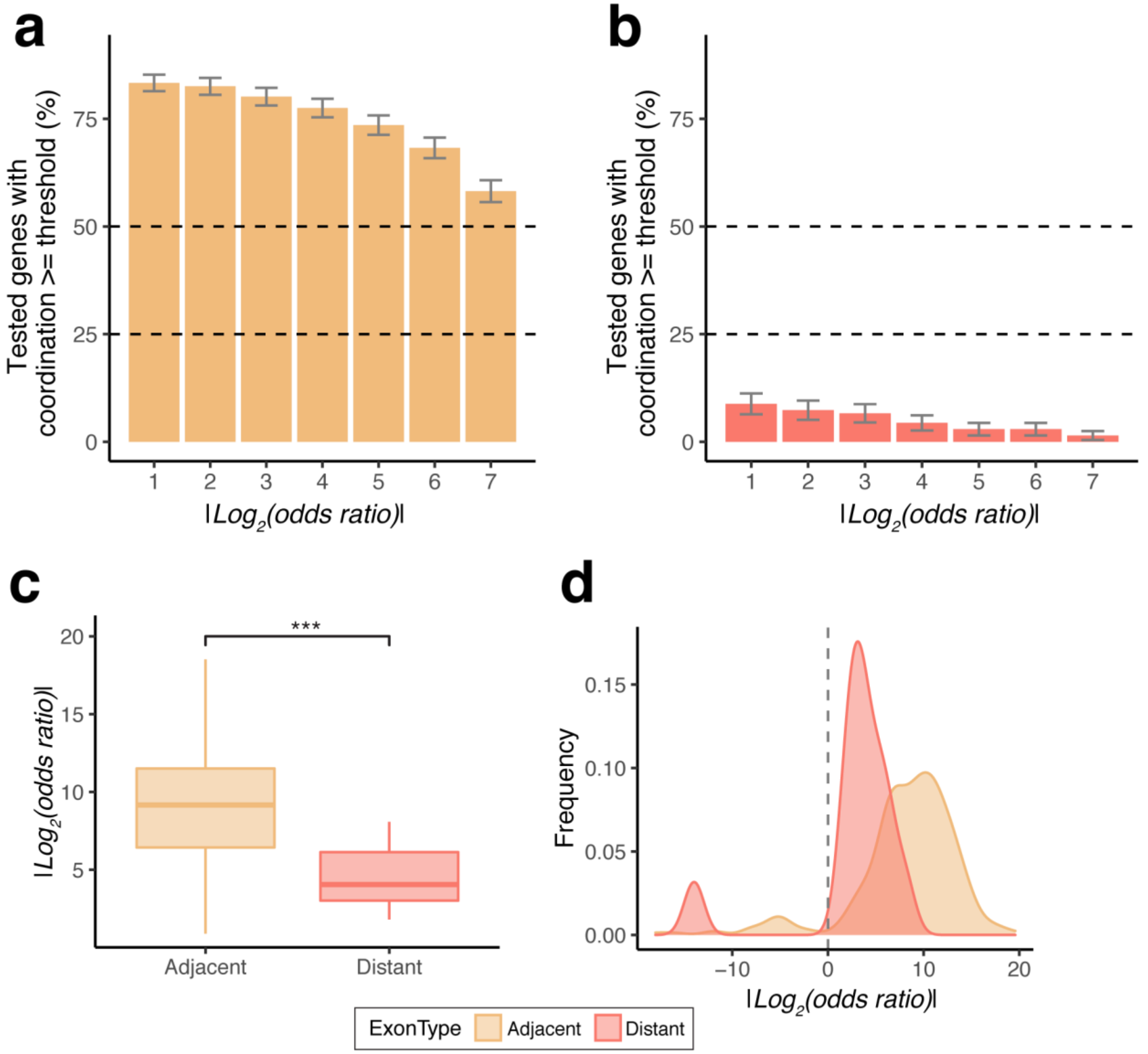


**Supplementary Figure 5: Replicable coordination of adjacent and distant exon pairs. a-b.** Barplot showing percent of tested genes in pseudo-bulk with significant exon coordination (FDR≤0.05) for adjacent (a) and distant (b) exon pairs at various log-odds ratio (LOR) cutoffs on the X-axis. Error bars indicate standard error of the point estimate. **c.** Boxplots of the absolute value of LOR for significant genes (FDR ≤ 0.05) on the Y-axis plotted against adjacent and distant exon pairs (A-B) on the X-axis. p-value obtained from two-sided Wilcoxon rank sum test. **d.** Density plot for the LOR for adjacent and distant exon pairs.

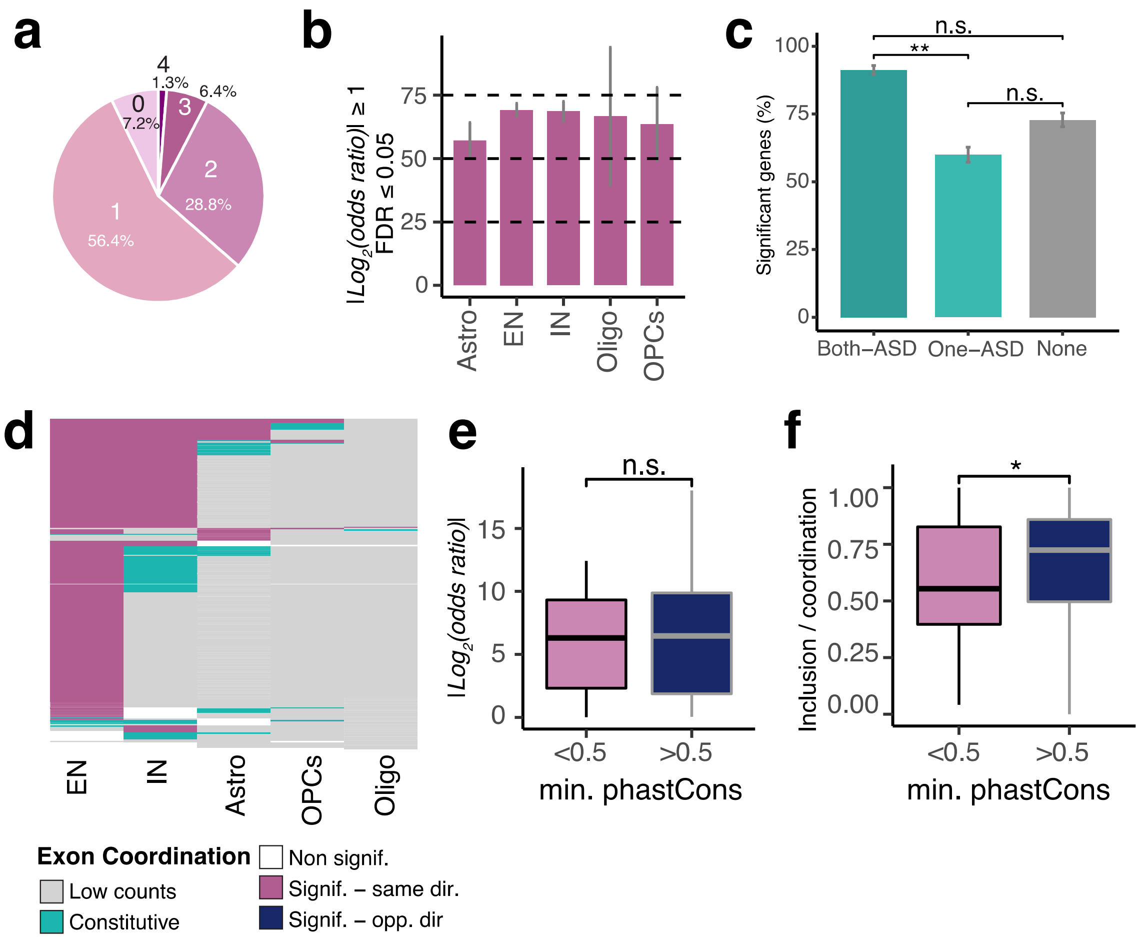


**Supplementary Figure 6: Exon coordination patterns across multiple cell types are replicable.** **a.** Pie chart indicating the number of cell types where an exon pair is significant (FDR ≤ 0.05 and abs(LOR)≥1) given that the exon-pair is testable in at least one cell type, and is significant when tested in pseudo-bulk. **b.** Barplot indicating percentage of tested exon-pairs (one per gene) that were deemed significantly coordinated (FDR ≤ 0.05 and abs(LOR) ≥ 1). Breakdown by cell type on the X-axis, given significance in pseudo-bulk. Error bars represent the standard error of the point estimate. **c.** Barplots showing percent of tested genes (FDR≤0.05, abs(LOR) ≥1) in pseudo-bulk for distant exons pairs on the X-axis where either one, or both, or neither of the exons are known to be associated with ASD. Error bars indicate standard error of the point estimate and p-values obtained from two-sided Fisher's exact test. **d.** Heatmap showing cell types as columns and exon pairs that were testable in at least one cell type as rows. Each element of the heatmap is colored by whether the exon-pair showed significant coordination and the direction of coordination with respect to the pseudo-bulk (same: pink or opposite: blue), was not significant (white), was not testable in a particular cell type because it had too few counts to satisfy the $\chi^2$ criterion (grey) or because one or both exons became constitutively included in a particular cell type (teal). **e.** Boxplots of the absolute value of the LOR for tested exon pairs for the excitatory neuron reads denoted on the Y-axis and split by whether the minimum phastCons score of the two exons was greater than or less than 0.5. Significance calculated using two-sided Wilcoxon rank sum test. **f.** Boxplots for the excitatory neuron reads showing the double inclusion level of an exon pair divided by the number of reads that were coordinated i.e. double inclusion + double exclusion. X-axis indicated whether the minimum phastCons score of the two exons was greater than or less than 0.5. Significance was calculated using the two-sided Wilcoxon rank sum test.

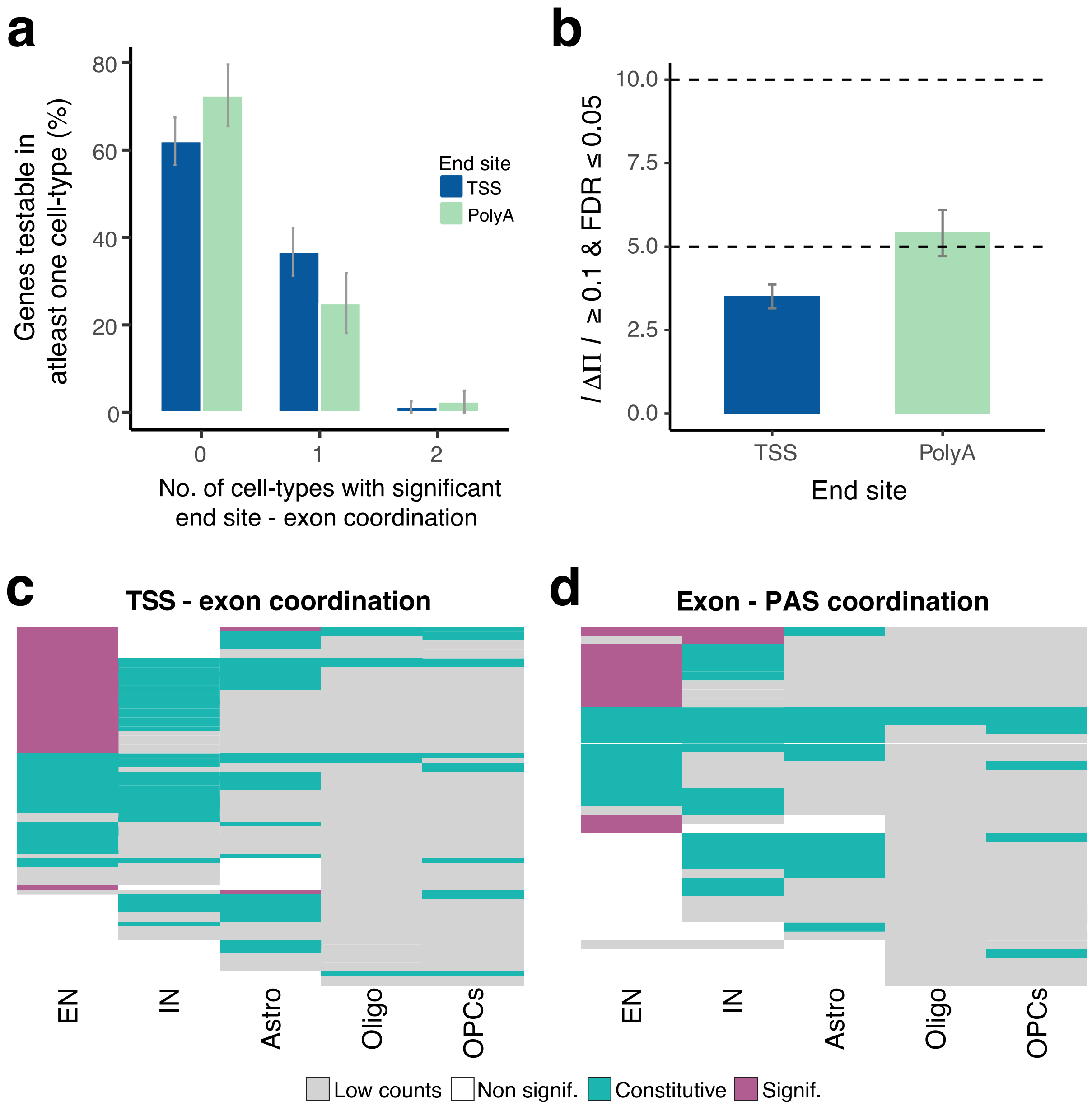


**Supplementary Figure 7: Cell type mediated exon - end site coordination in FCtx2. a.** Bar chart indicating the number of cell types where an exon-end site pair is significant (FDR ≤ 0.05 and **ΔΠ**≥1) given that the exon-end site pair is significant when tested in pseudo-bulk and is testable in at least one cell type. Color of bar indicates whether the end site associated with an exon is a TSS (blue) or PAS (green). Error bars represent standard error of the point estimate. **b.** Bar chart showing percent of tested genes in pseudo-bulk with significant exon-end site coordination (FDR ≤ 0.05) on the Y-axis and various ***ΔΠ*** cutoffs on the X-axis. Color of bar indicates whether the end site associated with an exon is a TSS (blue) or PAS (green). Error bars indicate standard error of the point estimate. **c-d.** Heatmaps showing cell types as columns and exon-end site pairs (exon-TSS left, exon-PAS right) that were testable in at least one cell type as rows. Each element of the heatmap is colored by whether the exon-end site pair showed significant coordination (pink), was not significant (white), was not testable in a particular cell type because it had too few counts to satisfy the $\chi^2$ criterion (grey), or because the exon or one or more end sites became constitutively included in a particular cell type (teal).

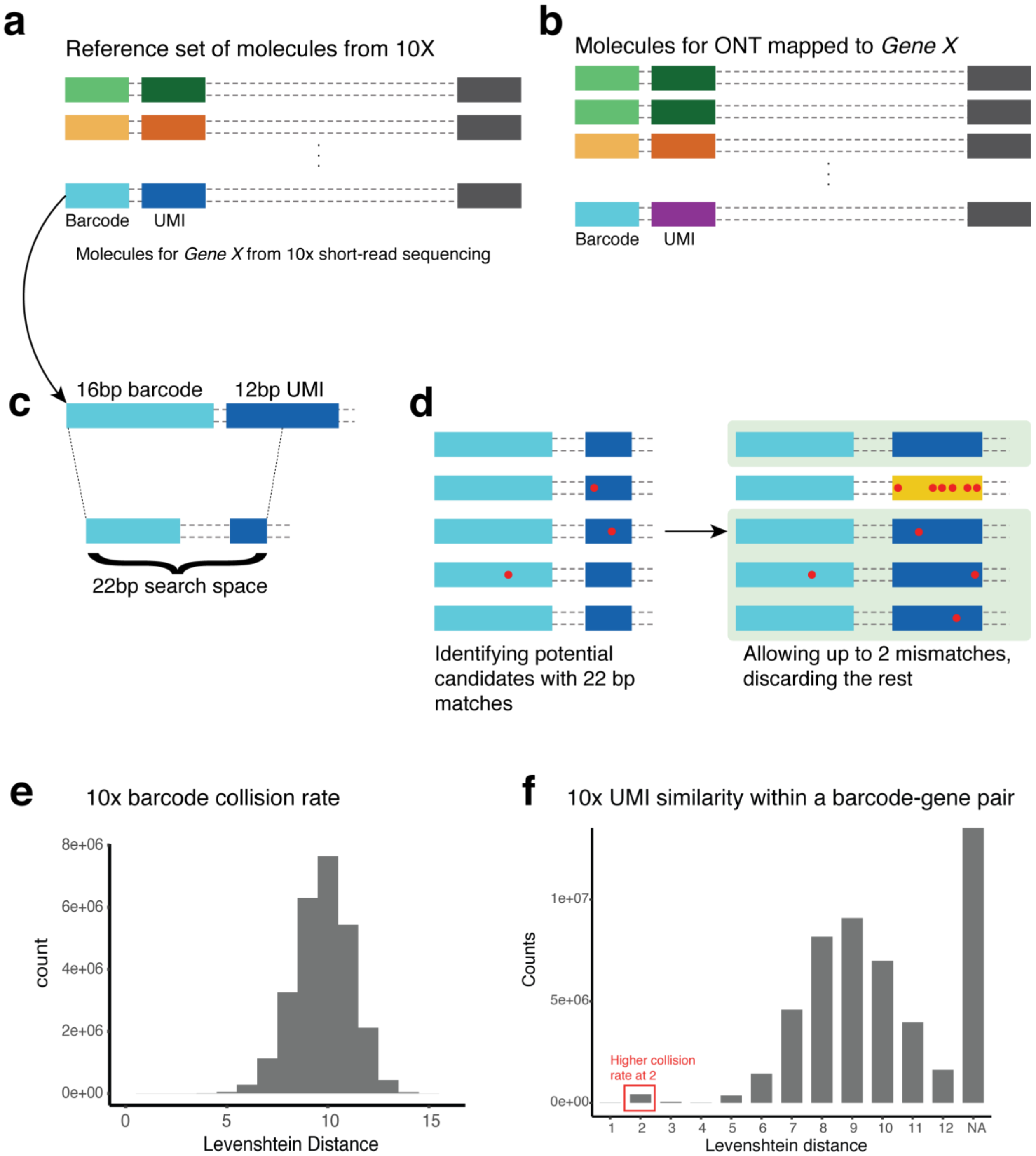


**Supplementary Figure 8: Barcode detection for long read transcripts obtained from ONT. a.** Reference set of molecules (Barcode-UMI sequence) for *GeneX* from 10x short-read sequencing data. **b.** ONT reads mapped to reference *GeneX.* **c.** First 22 bp of 28 bp long reference sequence used as a search space. **d.** Isolating ONT reads with up to 1 mismatch in 22 bp of reference sequence shown in C. Getting the full 28 bp barcode + UMI sequence and only retaining reads with up to 2 mismatches. **e.** Histogram of Levenshtein distances of every pairwise comparison of barcodes found in 10x FCtx1 sample. **f.** Histogram of Levenshtein distances of every pairwise comparison of UMIs per gene-barcode pair reported in 10x FCtx1 sample. Red box highlights a higher than expected rate for 2 mismatches within UMIs for a single gene-barcode pair.

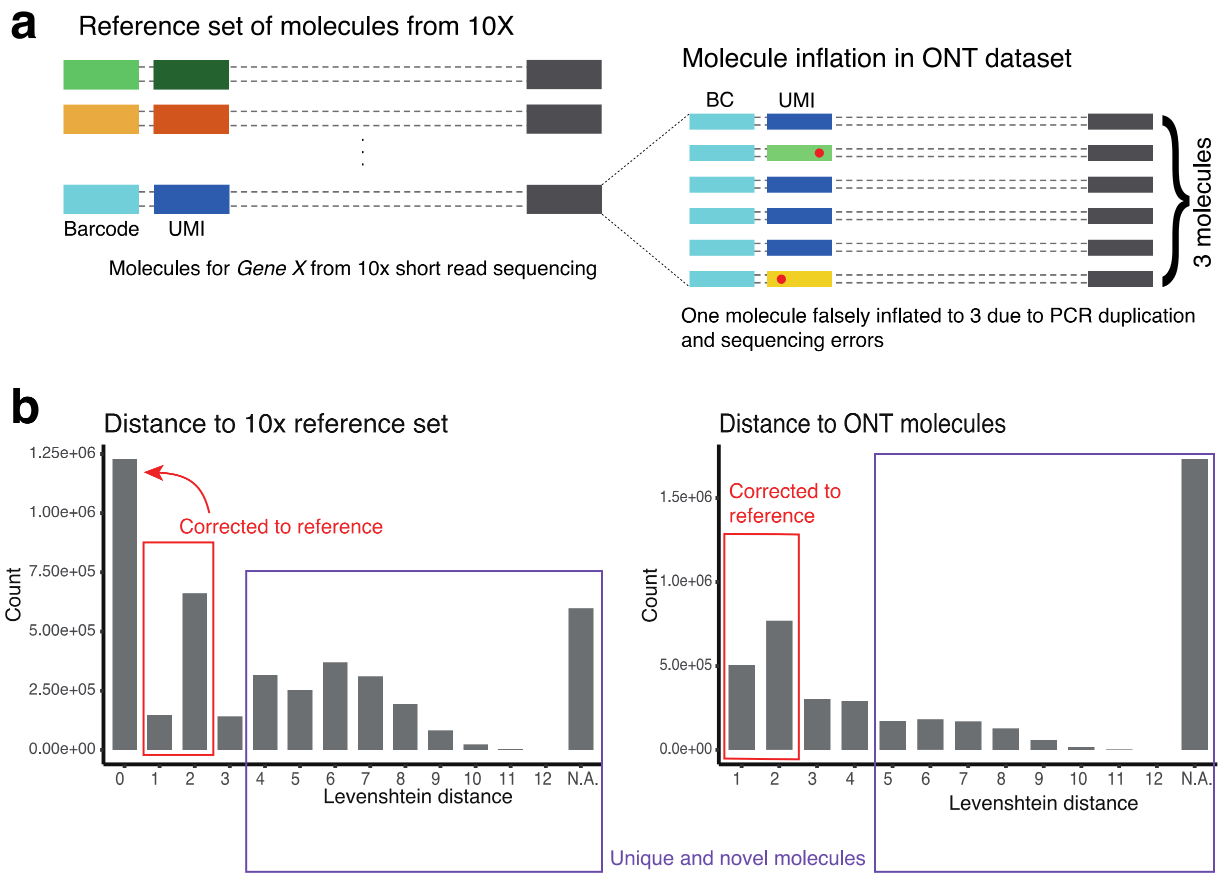


**Supplementary Figure 9: Identification of unique molecules from ONT data. a.** Illustration of 'molecule inflation' caused by PCR duplication and sequencing errors in ONT data for a single barcode-UMI pair associated with a gene. **b.** Histogram of Levenshtein distances of every pairwise comparisons of UMIs per gene-barcode pair found in the FCtx1 sample. *Left:* comparisons of ONT UMIs with the 10x reference set. *Right:* comparisons of ONT UMIs with other ONT UMIs associated with a gene-barcode pair. Red boxes highlight the fraction of UMIs that were corrected to the reference. Purple box highlights the UMIs which were sufficiently different from all existing UMIs and were therefore considered to be novel.

**Supplementary Table 1: Long-read sequencing statistics**

| Sample | Platform | Flow cell | No. reads | Mapped | On-target reads (%) | Avg. read length (bp) |
|---|---|---|---|---|---|---|
| FCtx1 (post-cap) | ONT | FCtx1_ONT_Run1.fastq.gz | 18,668,000 | 15,309,783 | 72.33 | 1283.6 |
| | | FCtx1_ONT_Run2.fastq.gz | 65,880,057 | 54,592,985 | 71.93 | 1157.7 |
| | | FCtx1_ONT_Run3.fastq.gz | 71,328,400 | 59,578,396 | 69.27 | 1123.3 |
| FCtx2 (post-cap) | ONT | FCtx2_ONT_Run1.fastq.gz | 44,079,800 | 38,466,551 | 68.81 | 864.2 |
| | | FCtx2_ONT_Run2.fastq.gz | 61,576,667 | 52,461,421 | 74.63 | 964.5 |
| FCtx1 (post-cap) | PacBio | FCtx1_PacBio_Run1.fastq.gz | 1,994,964 | 1,736,962 | 77.52 | 1112.1 |
| | | FCtx1_PacBio_Run2.fastq.gz | 2,913,665 | 2,536,474 | 77.45 | 1111.7 |
| | | FCtx1_PacBio_Run3.fastq.gz | 2,559,272 | 2,237,625 | 78.04 | 1140.1 |
| | | FCtx1_PacBio_Run4.fastq.gz | 3,219,346 | 2,803,921 | 77.59 | 1114.3 |
| | | FCtx1_PacBio_Run5.fastq.gz | 2,447,659 | 2,125,993 | 77.54 | 1104.6 |
| | | FCtx1_PacBio_Run6.fastq.gz | 2,334,405 | 2,025,196 | 77.46 | 1108.0 |
| | | FCtx1_PacBio_Run7.fastq.gz | 2,731,126 | 2,402,928 | 78.57 | 1149.4 |
| | | FCtx1_PacBio_Run8.fastq.gz | 2,013,024 | 1,766,104 | 78.37 | 1150.9 |
| FCtx2 (post-cap) | PacBio | FCtx2_PacBio_Run1.fastq.gz | 1,069,301 | 919,510 | 60.63 | 1098.6 |
| | | FCtx2_PacBio_Run2.fastq.gz | 297,808 | 256,278 | 61.00 | 1083.4 |
| | | FCtx2_PacBio_Run3.fastq.gz | 549,338 | 471,551 | 61.06 | 1069.0 |
| | | FCtx2_PacBio_Run4.fastq.gz | 402,627 | 346,060 | 61.00 | 1067.0 |

| | | | | | | |
|---|---|---|---|---|---|---|
| | | FCtx2_PacBio_Run5.fastq.gz | 1,557,832 | 1,335,563 | 61.56 | 1076.7 |
| | | FCtx2_PacBio_Run6.fastq.gz | 754,952 | 647,741 | 61.49 | 1088.4 |
| | | FCtx2_PacBio_Run7.fastq.gz | 194,487 | 163,288 | 77.01 | 934.5 |
| FCtx1 (pre-cap) | PacBio | FCtx1_PacBio_preCAP.fastq.gz | 1,121,497 | 933,889 | 23.49 | 1219.1 |
| FCtx2 (pre-cap) | PacBio | FCtx2_PacBio_preCAP.fastq.gz | 1,487,748 | 1,264,707 | 24.74 | 1233.1 |

ONT: Oxford Nanopore Technologies; PacBio: Pacific Biosciences; FCtx: Frontal cortex.
